# A polymer gel index-matched to water enables diverse applications in fluorescence microscopy

**DOI:** 10.1101/2020.10.04.324996

**Authors:** Xiaofei Han, Yijun Su, Hamilton White, Kate M. O’Neill, Nicole Y. Morgan, Ryan Christensen, Deepika Potarazu, Harshad D. Vishwasrao, Stephen Xu, Yilun Sun, Shar-yin Huang, Mark W. Moyle, Qionghai Dai, Yves Pommier, Edward Giniger, Dirk R. Albrecht, Roland Probst, Hari Shroff

## Abstract

We demonstrate diffraction-limited and super-resolution imaging through thick layers (tens-hundreds of microns) of BIO-133, a biocompatible, UV-curable, commercially available polymer with a refractive index (RI) matched to water. We show that cells can be directly grown on BIO-133 substrates without the need for surface passivation and use this capability to perform extended time-lapse volumetric imaging of cellular dynamics 1) at isotropic resolution using dual-view light-sheet microscopy, and 2) at super-resolution using instant structured illumination microscopy. BIO-133 also enables immobilization of 1) *Drosophila* tissue, allowing us to track membrane puncta in pioneer neurons, and 2) *Caenorhabditis elegans*, which allows us to image and inspect fine neural structure and to track pan-neuronal calcium activity over hundreds of volumes. Finally, BIO-133 is compatible with other microfluidic materials, enabling optical and chemical perturbation of immobilized samples, as we demonstrate by performing drug and optogenetic stimulation on cells and *C. elegans*.

## Introduction

Fluorescence microscopy spurs biological discovery, especially if imaging is performed at high spatiotemporal resolution and under physiologically relevant conditions. Coupling fluorescence microscopy with strategies for immobilizing or confining samples enables further applications, particularly when studying organisms that move rapidly. For example, the transparency and genetic accessibility^1^ of the nematode *C. elegans* has made it an ideal system for studying the growth, morphology and function of individual cells in the context of the whole organism^2,3^; yet imaging the living animal without motion blur usually requires immobilization with chemical^4,5^, steric^6,7^, or microfluidic^8-11^ means.

Microfluidic systems provide efficient immobilization and handling^12-18^ for studying cellular morphology^19,20^ and dynamics^21^, neuronal function^22-24^, behavior^25-27^, and lifespan^28,29^. Hydrogels (either independently^7,30^ or in conjunction with microfluidics^31^) have also been demonstrated as highly useful materials with tunable mechanical^32^, diffusive, and optical properties^33^ that are well-suited for long-term imaging applications^7,34,35^.

Unfortunately, relatively few attempts have been made to index-match immobilization devices^36-39^. The high refractive index (RI, n) of materials commonly used in microfluidics, such as polydimethoxylsilane (PDMS), causes significant optical aberrations^40,41^ due to the RI mismatch that occurs at the interface between the polymer (n_PDMS_ ∼ 1.41) and an aqueous sample (n_water_ = 1.33). These aberrations severely degrade image focus, resolution, and signal, compromising the performance of immobilization devices by reducing the information content of the resulting data. Hydrogels offer a lower RI (n ranging from 1.34 – 1.41) depending on thickness and polymerization conditions^33^. Although the RI of these materials is better matched to living samples, even a small mismatch in RI causes a noticeable deterioration in image quality^42^. Image degradation is particularly obvious when using water-dipping lenses designed for imaging living samples, such as those employed in high-resolution light-sheet fluorescence microscopy (LSFM)^43-45^.

Here we demonstrate a broadly applicable refractive-index-matched specimen mounting method that introduces negligible aberration when imaging living samples with high-resolution light-sheet microscopy and super-resolution microscopy. We show its utility in combination with microfluidics, enabling applications in high-resolution, volumetric imaging of cells, *Drosophila* tissue, and *C. elegans* adults and larvae. Our method takes advantage of the commercially available UV curable optical polymer BIO-133 (MY Polymers Ltd.) that has a refractive index matched to water (n = 1.333), is non-fluorescent, and is non-toxic. We show that 1) BIO-133 provides a gas permeable, inert, and biocompatible scaffold on which to grow and image tissue culture cells, 2) enables rapid tissue or animal encapsulation, and 3) is compatible with other microfluidic mounting schemes and optical or chemical perturbations.

## Results

### BIO-133 does not introduce additional optical aberrations

We assessed the optical properties of BIO-133 by using dual-view light-sheet microscopy (diSPIM^43,46^) to image 100 nm yellow-green beads placed under polymer layers of progressively increasing thickness (**Fig. 1, Methods**). In the most common diSPIM implementation, two identical 0.8 numerical aperture (NA) water-dipping objectives mounted above a planar substrate (usually a glass coverslip) alternately illuminate the sample with a light sheet and detect the resulting fluorescence. Since both illumination and detection planes are angled at ∼45 degrees with respect to the glass coverslip, imaging through a polymer gel with surface parallel to the coverslip will introduce significant lateral (‘x’ direction, **Fig. 1a**) and axial (‘z’ direction, **Fig. 1a**) aberrations if the gel’s refractive index differs from that of water.

**Fig. 1.**
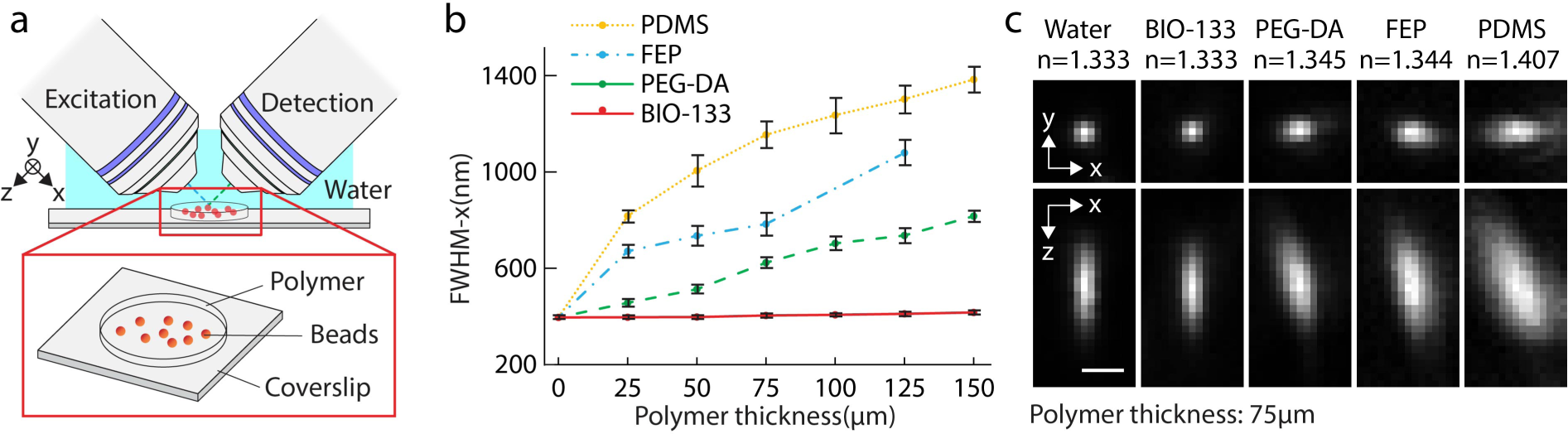
Diffraction-limited imaging is possible when imaging through BIO-133, unlike other polymers. **a)** Imaging geometry. A light sheet is used to illuminate the 100 nm yellow-green bead sample, which is embedded under progressively thicker polymer. Illumination and detection occur through 0.8 NA water-dipping objectives. **b)** Full width at half maximum (FWHM) in the ‘x’ direction under different thicknesses of polymer. Means and standard deviations are shown. **c)** Exemplary lateral (top row) and axial (bottom row) images of beads imaged through 75 μm of polymer, demonstrating that BIO-133 provides diffraction-limited performance whilst the other polymers do not. Single images, rather than maximum intensity projections, are shown. The refractive index of each polymer as measured with a refractometer is also indicated (average value from 3 independent trials). Scale bar: 1 μm. See also **Supplementary Table 1**.

As predicted, we observed this effect when imaging beads embedded under polymers with different RIs (**Fig. 1b, c**). Under no polymer, images of beads approximate the system point spread function, with measured lateral and axial full width at half maximum (FWHM) 395.9 +/- 7.7 nm and 1527.9 +/- 119.5 nm (*N* = 70 beads), respectively. Imaging beads under PDMS caused severe aberrations (**Fig. 1c**), more than doubling the lateral FWHM under a 25 μm layer (816.8 +/-24.9 nm) with progressive deterioration under thicker polymer layers (**Fig. 1b**). We also observed aberrations (**Fig. 1b, c**) under poly(ethylene-glycol) diacrylate (PEG-DA)^7^ and fluorinated ethylene polymer (FEP)^47^, albeit to lesser extent as the refractive indices of these polymers are closer to water. By contrast, beads imaged under BIO-133 showed negligible visual aberrations or measurable degradation in image quality (**Fig. 1b, c, Supplementary Table 1**), even under a 150 μm thick film, the largest thickness we tested (lateral FWHM 416.5 +/- 8.5 nm). We attribute this result to the refractive index of BIO-133, which we measured to be 1.333. We also verified, under 488 nm, 561 nm, and 637 nm illumination, that BIO-133 introduces negligible autofluorescence (**Supplementary Fig. 1**).

### High- and super-resolution imaging of cells through a layer of BIO-133

BIO-133 has permeability to oxygen about 2-3 times greater than PDMS and has water repellent properties (Personal Communication, Ehud Shchori, My Polymers, Ltd.). These material properties are advantageous for maintaining physiologically relevant cell culture conditions. Thus, we investigated whether BIO-133 could provide an inert and biocompatible scaffold for single cell imaging (**Supplementary Table 2a**). After a curing and leaching treatment (**Methods**), U2OS cells seeded on a 50 μm layer of BIO-133 adhered and displayed similar morphology and growth rate to cells grown on glass coverslips (**Fig. 2a, Supplementary Fig. 2**). Similar results were obtained using HCT-116 (human colon carcinoma) cells that express endogenous topoisomerase I-GFP, and we also observed similar expression and localization of tagged proteins compared to cells cultured on glass coverslips **(Supplementary Figs. 2, 3)**.

**Fig. 2.**
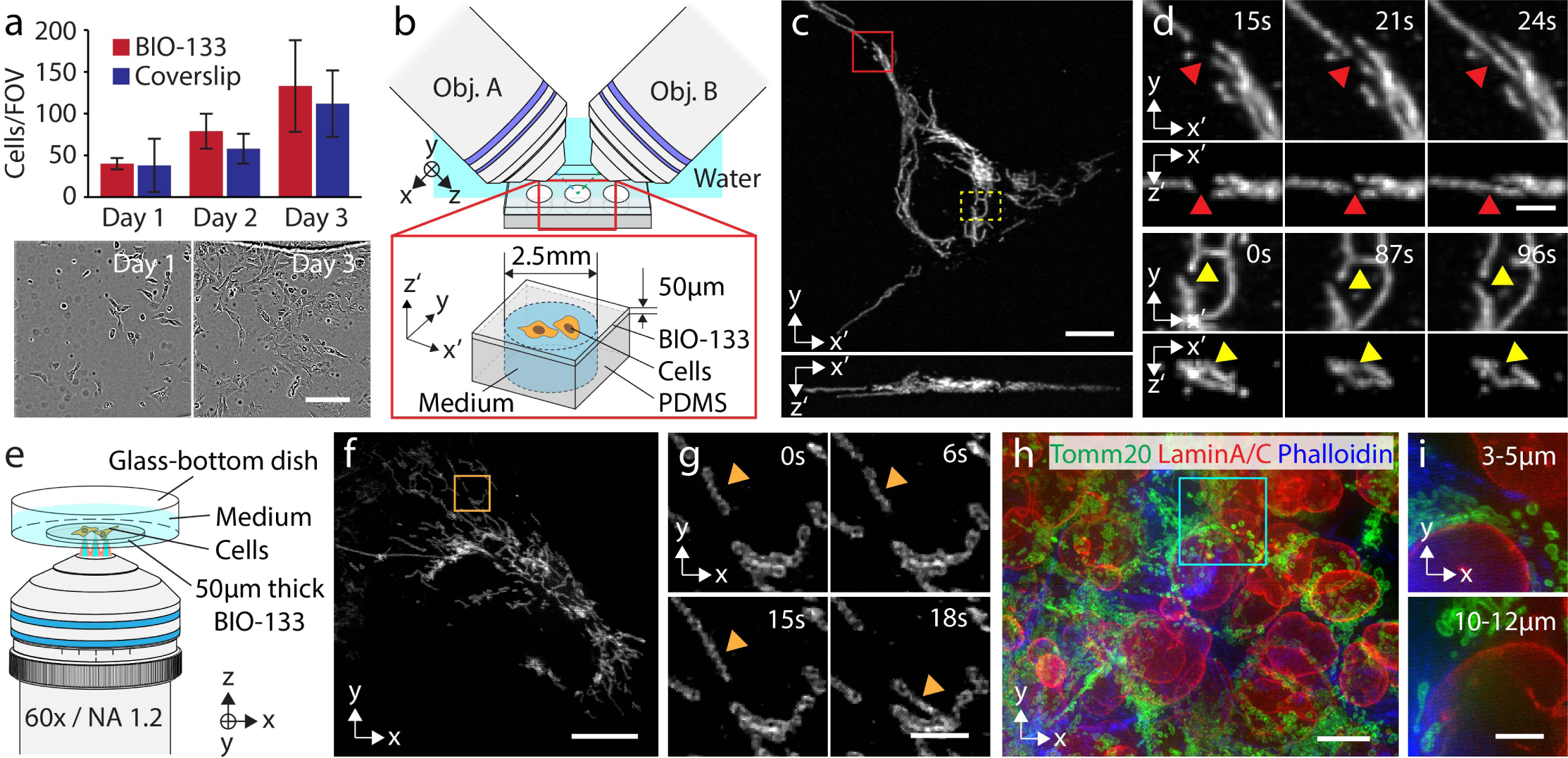
BIO-133 provides an inert and biocompatible scaffold on which to grow and image cells. **a)** U2OS growth on BIO-133 is similar to growth on glass coverslips. *Top*: quantifying cell growth on 50 μm thick BIO-133 layer vs. glass coverslip. Means and standard deviations from 3 fields of view (10x magnification, ∼800 μm x 800 μm field of view) are shown over 3 days. *Bottom*: example fields of view from day 1 and day 3, cells on BIO-133 layer. Scale bar: 200 μm. See also **Supplementary Fig. 2, 3. b)** Schematic of diSPIM imaging geometry. 50 μm film with adherent cells is inverted and imaged in the diSPIM setup. See also **Supplementary Fig. 4. c**) Example maximum intensity projections of deconvolved images of U2OS cells expressing mEmerald-Tomm20 in lateral (top) and axial (bottom) views. Scale bar: 10 μm. **d)**. Higher magnification views of the red and yellow rectangles in **c)**, highlighting examples of mitochondrial fusion (top, red arrowhead) and fission (bottom, yellow arrowhead). 50 volumes were taken with a 3 s inter-volume interval. See also **Supplementary Video 1**. Note that primed coordinates refer to the plane of the BIO-133 layer (x’, y) and the direction normal to the BIO-133 layer (z’). Scale bar: 2 μm. **e**) iSIM imaging geometry. Cells were cultured on 50 μm BIO-133 film, and the film placed in a glass-bottom dish and immersed in cell culture medium. Imaging was performed with a 60x, NA 1.2 water-immersion lens. **f**) Example deconvolved iSIM maximum intensity projection showing live U2OS cells expressing mEmerald-Tomm20. Scale bar: 10 μm. **g)** Higher magnification view of orange rectangular region in **f)**. Orange arrowhead marks the same mitochondrion. 25 volumes were acquired with a 3 s inter-volume interval. See also **Supplementary Videos, 3, 4**. Scale bar: 2 μm. A 0.5 pixel median filter was used to denoise images in **f, g)** prior to display. **h)** Multiple layers of HCT-116 cells were grown on 50 μm BIO-133 layer and immunostained against Tomm20 (green), lamin A/C (red), and actin (blue). See also **Supplementary Video 5**. Scale bar: 10 μm. **i)** Maximum intensity projection over indicated axial range (measured from the bottom of the cell layer) for cyan rectangular region in **h)**. Scale bar: 5 μm.

To demonstrate that transfected cells seeded directly on BIO-133 could be imaged at high spatiotemporal resolution, we created BIO-133 substrates on PDMS supports (**Supplementary Fig. 4**) and imaged cells expressing mEmerald-Tomm20, a fluorescent marker of the outer mitochondrial membrane, through a 50 μm thick BIO-133 layer using diSPIM (**Fig. 2b, c**). The jointly registered and deconvolved data acquired from two views displayed isotropic spatial resolution (**Fig. 2c, d**), allowing us to clearly visualize individual mitochondria and their dynamics (**Supplementary Video 1**), including mitochondrial fusion and fission (**Fig. 2d**). We also used BIO-133 in conjunction with diSPIM to construct a simple gravity-driven flow cytometry setup, obtaining clear images of DAPI stained nuclei as they flowed through the chamber (**Supplementary Fig. 5, 6, Supplementary Video 2**).

Next, we sought to image subcellular targets at spatial resolution beyond the diffraction limit, so we turned to instant structured illumination microscopy (iSIM)^48^, which enables high-speed super-resolution imaging. We again seeded U2OS cells expressing mEmerald-Tomm20 on a 50 μm BIO-133 film, this time imaging them with iSIM using a water dipping lens in an inverted geometry (**Fig. 2e**). Again, BIO-133 provided aberration-free imaging, enabling us to visualize the internal mitochondrial space absent Tomm20 (**Fig. 2f, g, Supplementary Video 3**). We also visualized LAMP-1-GFP-stained lysosome dynamics in wild type HCT-116 cells grown on another BIO-133 film of 50 μm thickness (**Supplementary Fig. 7, Supplementary Video 4**). As a third example, we grew multiple layers of HCT-116 cells on BIO-133, and immunostained the cells for lamin A/C, Tomm20, and actin (**Fig. 2h**), obtaining clear images of these structures through the volume of the sample (**Fig. 2i, Supplementary Video 5**). We conclude that BIO-133 is compatible with multicolor, super-resolution imaging in live and fixed targets.

### BIO-133 enables subcellular imaging, segmentation, and tracking within immobilized living tissue

In addition to monitoring the dynamics of organelles within single cells, we also immobilized and imaged multicellular structures in flies and worms at high spatiotemporal resolution, using diSPIM in conjunction with BIO-133 for sample immobilization (**Fig. 3, Supplementary Table 2b**). The developing *Drosophila* wing has long been a model for axon growth and neuronal pathfinding and differentiation^49,50^. More recently, spinning disk confocal microscopy was used to dissect the role of cytoskeletal organization and dynamics in shaping the morphogenesis and growth of the TSM1 pioneer sensory neuron axon in explanted early-pupal wing imaginal discs^51,52^. In those experiments, phototoxicity and photobleaching limited imaging duration to ∼30 volumes, with image volumes acquired every 3 minutes. Wings were sandwiched between two glass coverslips to immobilize the preparation and prevent it from moving during imaging, but this scheme introduces unacceptable aberrations if imaging with the less phototoxic diSPIM. Instead, we immobilized wings with a thin layer of BIO-133 (**Fig. 3a, Supplementary Fig. 8**), which enabled sustained volumetric imaging with diSPIM. We acquired 360 single-view volumes (5 s inter-volume interval, spanning 30 minutes) of tdTomato-CD4 expressed in TSM1 and the neighboring L3 neuron, marking neuronal membranes (**Fig. 3b**). In addition to observing slower remodeling of the TSM1 growth cone (**Fig. 3c, top**) our imaging rate also enabled us to capture rapid movement of membrane-labeled puncta that appeared to traffic along the L3 axon shaft (**Fig. 3c, bottom, Supplementary Video 6**). Puncta were also evident in comparative spinning disk confocal datasets (**Supplementary Fig. 9**).

**Fig. 3.**
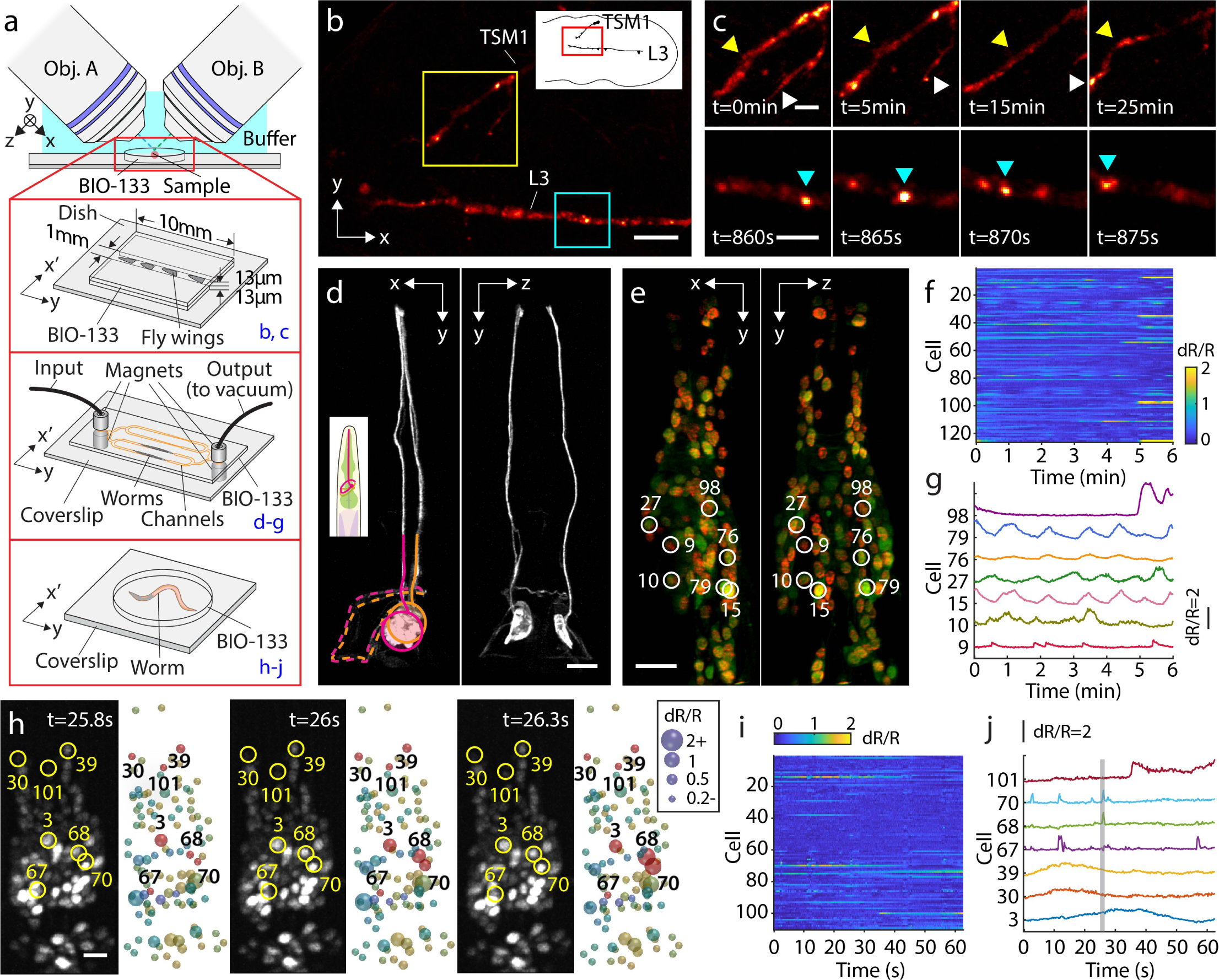
Live imaging of BIO-133 encapsulated fly wings and *C. elegans*. **a)** Experimental schematic. Top: A thin layer of BIO-133 membrane covers excised fly wings, immobilizing them so that axon dynamics can be recorded at high resolution over an extended period. Middle: A simple microfluidic device is used to trap adult worms for structural and functional imaging of the nervous system. Bottom: Worms can also be encapsulated in a gelled droplet of BIO-133. See also **Supplementary Figs. 8, 10. b)** Deconvolved, single-view maximum intensity projection of a fly wing with TdTomato-labelled CD4 showing the axons of two neurons (upper: TSM1; lower: L3) shortly before fasciculation in the developing *Drosophila melanogaster* wing disc. Scale bar: 10 μm. 360 volumes were taken with 5 s inter-volume intervals. (30 min in total, see also **Supplementary Video 6**). **c)** Magnified regions of TSM1 and L3 axons, corresponding to yellow and blue rectangles in **b)**, highlighting morphological changes and apparent motion of CD4 puncta. Scale bars: 4 μm. **d)** Isotropic, high-resolution imaging of GFP-labeled axons and dendrites in anesthetized adult *C. elegans*, as shown by orthogonal, jointly deconvolved diSPIM maximum intensity projections. Cell bodies are circled and axons entering the nerve ring region are overlaid with dotted lines. Scale bar: 10 μm. **e)** Calcium imaging of adult worm (red channel: TagRFP, green channel: GCaMP6s; both labels targeted to nuclei), 2 views imaged at 1.25 Hz volumetric rate. Joint deconvolution diSPIM results are shown; red and green channels were simultaneously collected and colors are overlaid in display. Scalebar: 10 μm. See also **Supplementary Video 7. f)** dR/R traces for all 126 tracked nuclei. **g)** dR/R traces for selected individual neurons. Note correspondence with numbered neurons and marked neurons in **e). h)** Calcium imaging of larval worm with higher temporal resolution (4 Hz volumetric rate), single-view results are shown. GCaMP channel and associated segmented dR/R signal are indicated for 3 successive time points. Scale bar: 5 μm. See also **Supplementary Video 8. i)** dR/R traces for all 110 nuclei segmented and tracked in **h). j)** traces for selected individual neurons. Note correspondence with numbered neurons and marked neurons in **h)**.

In another example, we used soft lithography techniques^53^ to cast BIO-133 into microfluidic devices suitable for trapping *C. elegans* (**Supplementary Fig. 10**). Introducing adult worms into the channels via suction **(Fig. 3a**), we imaged fine structures (**Fig. 3d**) and functional activity **(Fig. 3e-j**) in living animals at isotropic resolution. Animals were sufficiently immobile that we could serially acquire and fuse the two diSPIM views^54^ to obtain reconstructions free of motion blur. In strains expressing GFP sparsely targeted to a few neurons, we resolved axons and dendrites (likely from amphid neurons, **Fig. 3d**) within anesthetized *C. elegans*. When imaging the genetically encoded calcium indicator GCaMP6s^55^ and mCherry targeted pan-neuronally^56^ in immobilized adult animals without anesthetic, our volume imaging rate of 1.25 Hz (simultaneous acquisition of red and green channels) enabled us to segment and track 126 nuclei in the animal head (**Fig. 3e**,**f Supplementary Video 7**), permitting inspection of spontaneously active nuclei (**Fig. 3f, g**) over our 450 volume (6 minute) experiment. Intriguingly, we observed a pair of nuclei (#79 and #15, **Fig. 3e, g**) that exhibited in-phase, rhythmic activity with slow (45-80 s) period (**Supplementary Video 7**), as well as nuclei showing out-of-phase activity with respect to this pair (#27). In another experiment, we simply embedded *C. elegans* larvae expressing the same pan-nuclear GCaMP6s marker in a cured disk of BIO-133 (**Fig. 3a**), recording volumes from one side to obtain volumes at 4 Hz, for 250 volumes. Despite the poorer axial resolution of single-sided imaging, and the smaller size of the larval nuclei, we were able to again segment and track 110 nuclei in the head of the animal, identifying calcium transients in spontaneously active nuclei with a time resolution of 0.25 s (**Fig. 3h-j, Supplementary Video 8**).

The droplet-based design also enabled easy recovery of animals post-imaging. 26 / 28 animals were recovered even ∼12 hours after embedding, confirming our suspicion that cured BIO-133 is inert, gas permeable, water repellant, and does not obviously affect animal viability (the remaining two animals died within the BIO-133 capsule due to internal hatching of embryos within the animals). The water repellency of BIO-133 likely contributes to retaining the animal’s intrinsic hydration and thus viability during encapsulation. The ease at which *C. elegans* can be immobilized and imaged at high spatiotemporal resolution suggests useful synergy with multicolor strategies that permit unambiguous neural identification^23^.

### BIO-133 is compatible with chemical and optogenetic perturbations

The ability to specifically perturb and subsequently follow biological processes by observing morphological or functional changes is valuable in dissecting biological processes. We conducted several studies to show that BIO-133-mounted samples are compatible with such perturbations (**Fig. 4**). First, we conducted a simple drug assay by modifying our BIO-133/PDMS cellular scaffolds (**Fig. 2b, Supplementary Fig. 11**) so that U2OS cells could be exposed (**Fig. 4a**) to carbonyl cyanide m-chlorophenyl hydrazine (CCCP), an inhibitor of oxidative phosphorylation. Because we could clearly observe cells through the BIO-133 layer using diSPIM, we observed that, compared to control cells in a neighboring well (**Fig. 4a-c**), within minutes of exposure the treated cells showed mitochondrial fragmentation, eventually exhibiting major disruption to the mitochondrial network (**Fig. 4b, c, Supplementary Video 9**).

**Fig. 4.**
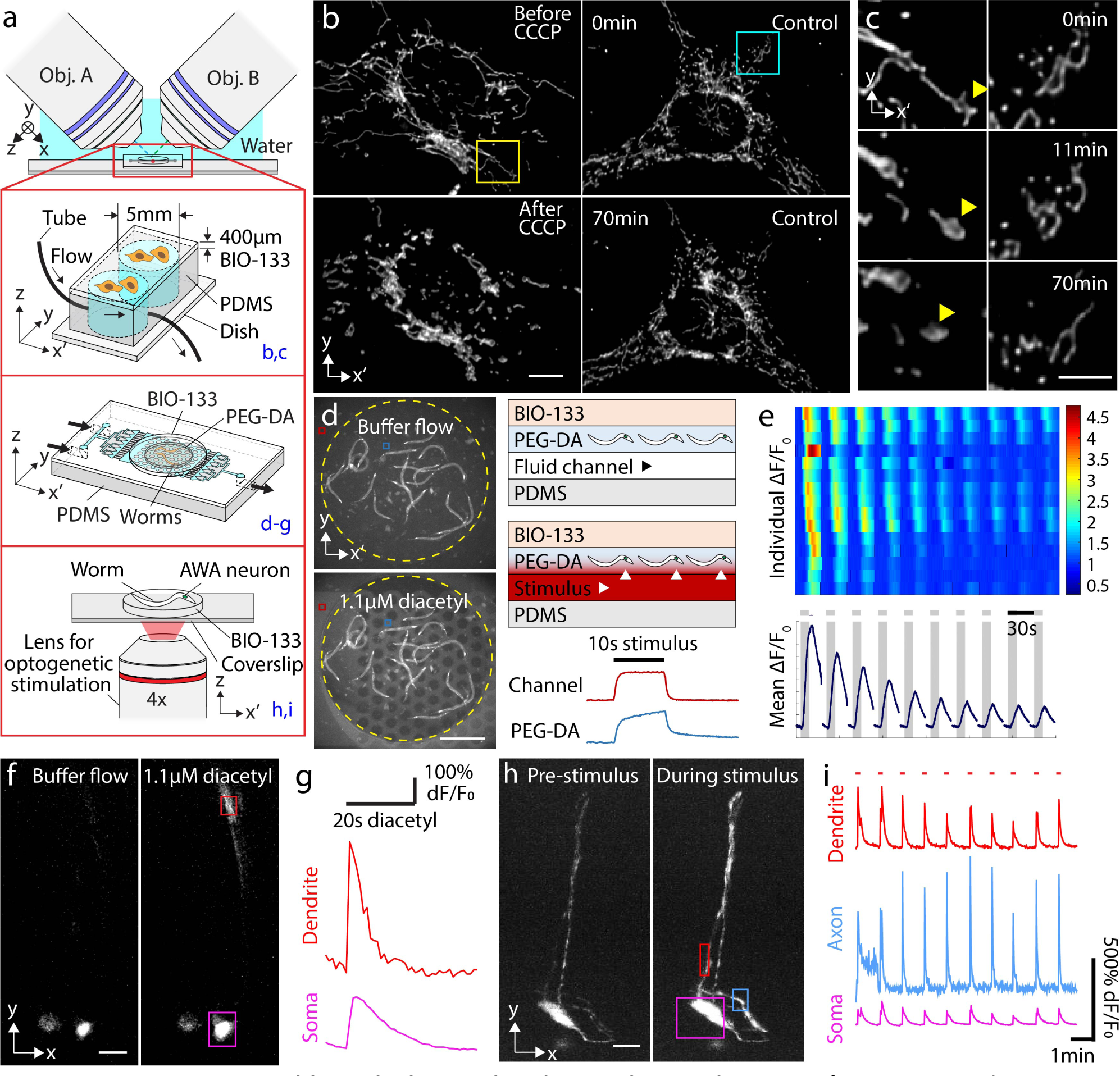
BIO-133 is compatible with chemical and optical perturbations. **a)** Experimental schematic for perturbations. Cells grown on BIO-133 were placed on PDMS wells and were either perturbed by flowing 0.05 mM CCCP or left as controls (top higher magnification view). Alternatively, worms were embedded in PEG-DA bonded to a PDMS flow chip and imaged through a layer of BIO-133 (middle higher magnification view) to examine response to chemical stimulation; or embedded in BIO-133, repetitively stimulated with red light from lower, 4x objective and imaged using upper diSPIM objectives (bottom higher magnification view). See also **Supplementary Figs. 11-13. b)** Example cells with (left column) and without (right column) CCCP treatment at early (top, 0 min) and late (bottom, 70 min) time points. CCCP was added at 10 minutes. Maximum intensity projections of deconvolved diSPIM data are shown. Scale bar: 10 μm. See also **Supplementary Video 9. c)** Higher magnification views of yellow (left column) and blue (right column) regions in **b)**. Yellow arrowhead shows CCCP-induced morphological change of mitochondrion. Scale bar: 5 μm. **d)** Example images of worms expressing GCaMP immobilized in PEG-DA disk with (bottom) and without (top) 1.1 μM diacetyl. Fluorescein added to stimulus highlights the rapid addition/removal of chemical. Scale bar: 500 μm. Right schematics show layered structure of assembly, including direction of flow and diffusion (arrowheads) into PEG-DA layer. Line plots show intensity of fluorescein over time in channel (red) and PEG-DA (blue) layers. **e)** Top: dF/F heatmaps derived from widefield microscopy measurements from 15 animals (rows) in response to 10 repeated stimulus pulses (once per minute). Bottom: responses averaged over all animals show neural adaptation. **f**) Single-view diSPIM images recorded from a single animal, showing subcellular response in AWA neuron to 1.1 μM diacetyl compared to control (buffer flow) conditions. Contrast has been adjusted to better highlight the response from different cell regions. Scale bar: 10 μm. See also **Supplementary Video 10. g)** Graphs show average intensity from boxed regions in **f)** highlighting fluorescence intensity changes in soma and dendrite. **h)** Worms expressing Chrimson and GCaMP are repetitively stimulated with red light and imaged using upper diSPIM objectives. Maximum intensity projection of GCaMP fluorescence from single-view diSPIM recordings are shown before (left) and after (right) optogenetic stimulation. Scale bar: 10 μm. See also **Supplementary Video 11. i)** dF/F traces for dendrite, axon, and soma, corresponding to boxed regions in **h)**.

Chemically stimulating animals directly embedded in BIO-133 is difficult, since BIO-133 is not permeable to aqueous solutions. One solution is to introduce chemicals via microfluidic channels (such as those shown in **Fig. 3d-g**). Alternatively, we explored using PEG-DA for immobilization and aqueous permeability, above a microfluidic layer for stimulus introduction and control, and beneath a BIO-133 layer to enclose the fluidic path. Using a thin PEG-DA disk allows easy transfer of different embedded organisms on the same imaging setup, and repeated imaging of the same animals over many hours if desired. We constructed a hybrid multi-material device composed of a PDMS microfluidic base bonded to a BIO-133 upper membrane that sealed in a small PEG-DA disk containing tens of embedded nematodes^7^ (**Supplementary Fig. 12, Fig. 4a**). Chemicals applied via flow channels diffuse into the PEG-DA disk, evoking neural responses that can be imaged through the BIO-133 viewing layer with widefield microscopy or diSPIM (**Fig. 4d-g, Methods**). We embedded 15 animals expressing GCaMP2.2b in AWA chemosensory neurons^57^ in a PEG-DA disk and applied 1.1 μM diacetyl pulses, which directly activate these neurons via the ODR-10 chemoreceptor^57,58^. Using widefield microscopy, we recorded robust calcium transients from all animals and observed a characteristic sensory adaptation to repeated stimulation (**Fig. 4e**). The initiation of sensory neural responses varied slightly due to diffusion of diacetyl stimulus through the PEG-DA disk to animals embedded in different planes (**Supplementary Fig. 13**). Next, we examined individual neuron responses by using the same apparatus with diSPIM (**Fig. 4f**,**g Supplementary Video 10**). Our imaging provided sufficient spatial resolution to distinguish subcellular responses, observing faster on and off dynamics of fluorescent transients in the dendrites than in the soma^59^.

Other stimulation modalities (such as light, temperature, and mechanical vibration) are directly transmissible through BIO-133, and these can be applied directly to cells and organisms embedded in the polymer in a simpler preparation. Optogenetic neural activation is a particularly advantageous tool, allowing remote light-induced activation or suppression of neurons. We embedded nematodes expressing the red light activated cation channel Chrimson^60^ and GCaMP2.2b^7^ directly in BIO-133 disks and monitored calcium readout with diSPIM during repeated red light stimulation pulses (**Fig. 4a**,**h**,**i, Supplementary Video 11**). We observed increases in fluorescence after each stimulus (**Fig. 4h**), and again could clearly localize such transients to subcellular areas including soma, dendrite, and axon (**Fig. 4i**).

## Discussion

BIO-133 is commercially available, rapidly curing, gas permeable, inert, water repellent, and biocompatible. It is not autofluorescent under visible illumination and does not introduce additional aberration when imaging with water-dipping or water-immersion objective lenses designed for aqueous specimens. These characteristics make it well-suited to microfluidic experiments under physiological conditions, particularly with the many LSFM systems that use such lenses. We suspect that capillary mounting^47^, often used for mounting zebrafish in LSFM, could be improved if BIO-133 were used instead of the FEP material commonly used in this application. Our finding that cells can be directly grown on BIO-133 without additional surface treatment may prove useful in non-standard LSFM geometries that previously employed RI-matched materials with passivated surfaces^45^, or in ultra-high-throughput light-sheet imaging^61,62^.

We also found that BIO-133 does not noticeably degrade imaging performed in more traditional inverted microscope geometries, including in super-resolution imaging. This capability suggests that using BIO-133 could improve imaging in studies of cell morphology, mechanics, migration, and motility, e.g., when using micropillars^20,63^ or in traction force microscopy^64^.

We bonded BIO-133 to glass and PDMS with silicone-based adhesive tape, or reversibly to glass via van der Waals forces. The elastic modulus of cured BIO-133 (5 MPa) is similar to that of PDMS (3.7 MPa). Thus, similar to PDMS^65^, BIO-133 conforms to minor imperfections in glass and bonds to it by weak van der Waal forces, creating a reversible bond and a watertight seal. We suspect further tuning of adhesive, optical and mechanical properties of this intriguing polymer is possible but will depend on knowing the chemical formula, which is currently proprietary.

## Author Contributions

Conceived project: R.P., H.S. Designed experiments: X.H., Y.S., H.W., K.M.O., H.D.V., E.G., D.A., R.P., H.S. Provided technical advice and resources for microfluidics: N.M., R.P., H.W., D.A. Provided biological advice: R.C., Y.S., S.H., Y.P. Created new reagents: S.H., Y.P, M.W.M. Performed experiments: X.H., Y.S., H.W., K.M.O., D.P., H.D.V., R.P. Tracked nuclei in GCaMP imaging experiments: X.H., Y.S., S.X. Wrote paper with input from all authors: X.H., Y.S., H.W., D.A., R.P., H.S. Supervised research: Q.D., Y.P., E.G., D.A., R.P., H.S. Directed research: H.S.

## Acknowledgements

This research was funded in part by the National Institute of Biomedical Imaging and Bioengineering, the National Institute of Neurological Disorders and Stroke, and the Center for Cancer Research of the National Cancer Institute within the National Institutes of Health (Z01 BC 006161), and the National Science Foundation (CBET 1605679). We thank George Patterson for the use of his cell culture facilities, Leighton Duncan and Daniel Colón-Ramos for kindly providing strains and for conducting initial pilot experiments and their careful read, Evan Ardiel and Eviatar Yemini for providing helpful feedback on pan-nuclear GCaMP recordings, Ron Zohar and Ehud Shchori for providing useful information on BIO-133, and Hank Eden for providing helpful feedback on the manuscript. K.M.O. and E.G. were supported by NINDS Z01-NS003013 to E.G. K.M.O. was also jointly supported by AFOSR grant number FA9550-16-1-0052 to W. Losert at UMD College Park. M.W.M was supported by NIH grant F32-NS098616.

## Disclaimer

The NIH, its officers, and staff do not recommend or endorse any company, product, or service.

## Methods

### Sample preparation

#### U2OS, wild type (WT) HCT-116, and HCT-116 TOP1-GFP cell culture

U2OS (ATCC, HTB-96), WT HCT-116 (ATCC, CCL-247), and HCT-116 TOP1-GFP (see below) cells were cultured in DMEM (Lonza, 12-604F) media with 10% Fetal Bovine Serum at 37°C and 5% CO_2_.

To tag the genomic topoisomerase I (TOP1) in WT HCT-116 cells, sequence CCTCACTTGCCCTCGTGCCT targeting a CRISPR site 77nt after the stop codon of TOP1 was cloned into pX330. Homology arms (of ∼1 kb) upstream and downstream of the target site were cloned to flank a blasticidin-resistance gene, where the upstream homology arm was modified to replace the stop codon with a GFP domain connected to the protein-coding region of the last exon of TOP1 via a short poly-lysine linker. Both constructs were co-transfected into WT-HCT116 cell, followed by selection with 5 μg/mL of blasticidin 48 hours post transfection. GFP-positive cells were further selected by FACS.

#### Transfection of cells

Cells were cultured to 50% confluency and transfected using xTreme Gene HP DNA Transfection Reagent (Sigma, 6366236001). The transfection mixture contained 100 μL 1X PBS, 2 μL Transfection Reagent, and 200-1000 ng plasmid DNA. Cells were imaged 24-48 hours after transfection.

#### C. elegans samples

Nematode strains were grown on NGM plates seeded with OP50 bacteria. *C. elegans* imaged as young adults were synchronized by picking L4 stage worms 24 hours prior to the experiment and transferring them to seeded plates, and *C. elegans* imaged as larvae were directly picked from plates. Strain DCR6268 (*olaEx3632[pttx-3b::SL2::Pleckstrin homology domain::GFP::unc-54 3’UTR + pelt-7::mCh::NLS::unc-54 3’UTR]*) was used for imaging axons and dendrites (**Fig. 3d**). olaEx3632 was made by injecting plasmid DACR2285 (pttx-3b::SL2::Pleckstrin homology domain::GFP::unc-54 3’UTR) at 25 ng/uL and DACR2436 (pelt-7::mCh::NLS::unc-54 3’UTR) at 10 ng/μL. Strain AML32^56^ (*wtfIs5 [rab-3p::NLS::GCaMP6s + rab-3p::NLS::tagRFP]*) was used for pannuclear neuronal calcium imaging (**Fig. 3e-j**); strain NZ1091(*kyIs587 [gpa-6p::GCaMP2*.*2b; unc-122p::dsRed]; kyIs5662 [odr-7p::Chrimson:SL2:mCherry; elt-2p::mCherry]*^*7*^) was used for chemical (**Fig. 4d-g**) and optogenetic stimulation (**Fig. 4h, i**). For optogenetic stimulation, L4 stage animals were transferred to agar plates seeded with 62.5 μM all trans-retinal (ATR, Sigma Aldrich, R2500) for over 12 hours.

#### Drosophila samples

*Drosophila* stocks were obtained from the Bloomington *Drosophila* Stock Center: *neur-GAL4* (BL6393) and *UAS-CD4-td-Tomato* (BL35837). White prepupae were selected and aged for 13-14 h at 18°C followed by 1 h at 25°C (equivalent to 7.5–8 h at 25°C). These aged pupae were dissected in fresh culture media (CM; Schneider’s Drosophila media + 10% fetal bovine serum, both from Life Technologies), and wing discs were isolated from the aged pupae and stored in fresh CM prior to mounting with BIO-133.

### Characterizing the optical properties of polymers

#### Characterization of aberrations by visualizing fluorescent beads under different polymer layers

100 nm diameter yellow-green fluorescent beads (Invitrogen, F8803, 1:10000 dilution in water) were coated on #1.5 coverslips (24 mm x 50 mm, VWR, 48393-241) coated with 0.1% w/v poly-L-lysine (Sigma, P8920-100ML). We placed spacers (Precision Brand, 44910) of variable thickness on one of the coverslips and deposited droplets of BIO-133, 10% PEG-DA (ESIBIO, GS705) or UV-curable PDMS (Shin-Etsu Chemical, KER-4690) on the beads. The droplet was then covered with another coverslip coated with beads and compressed with an iron ring. BIO-133 and PEG-DA hydrogels were crosslinked at 312 nm (Spectroline ENB-280C) for 2 minutes. PDMS was crosslinked at 312 nm for 5 minutes and post-cured at room temperature for one day. Once cured, we separated the two cover glasses and kept the one with polymer/hydrogel on it. For imaging through FEP films of different thickness (CS Hyde, 23-1FEP-24, 25 μm; 23-2FEP-24, 50 μm; 23-3FEP-24, 75 μm; 23-5FEP-24, 125 μm), we immersed beads in 1μL water and covered the sample with the FEP film. All bead images were acquired with a symmetric 0.8/0.8 NA diSPIM^43^.

#### Measurement of refractive index of polymers

A refractometer (American Optical) was used to measure the refractive index of pure water, BIO-133 film (My Polymer, BIO-133, 25 μm), PDMS film (Shin-Etsu Chemical, KER-4690, 25 μm), FEP film (CS Hyde, 23-1FEP-24, 25 μm), and 10% PEG-DA hydrogel (100 μm). The refractive index for each material was measured 3 times and the average value reported in **Fig. 1**.

#### Measurement of BIO-133 autofluorescence

A 50 μm thick BIO-133 film was deposited on a glass bottom dish (MatTek, P35G-1.5-14-C). Images were acquired both on the BIO-133 area and an area without BIO-133, using an instant structured illumination microscope (iSIM^48^) with 40 ms exposure time and 45 mW 488 nm excitation, 70 mW 561 nm excitation, or 90 mW 639 nm excitation (measured with a power meter immediately prior to the objective). Care was taken to ensure the illumination was focused within the BIO-133 film. The two images were subtracted to measure the autofluorescence of BIO-133 relative to glass (**Supplementary Fig. 1**).

### Cell growth and imaging using BIO-133 substrates

#### Fabrication of BIO-133 cell culture wells for diSPIM experiments

BIO-133-sided PDMS substrates with 2.5 mm diameter wells (**Supplementary Fig. 4**) were used for live cell imaging experiments (**Fig. 2b-d**). To make the BIO-133 bottom, a BIO-133 droplet was positioned on a #1.5 glass coverslip (24 mm x 50 mm, VWR 48393-241) between two 50 μm plastic spacers (Precision Brand, 44910), covered with another coverslip, compressed with a glass slide (Ted Pella, 260386) and cured with a UV lamp (365nm, Spectroline ENB-280C) for 15 minutes. After curing, the BIO-133 film was peeled off and exposed to UV light for another 2 hours in 70% ethanol. To make the PDMS well, 15 mL PDMS (Dow Inc. Sylgard 184) was poured into a 10 cm plastic dish (Kord-Valmark, 2910) and cured for 2 hours at 80 °C to obtain a 2 mm thick PDMS slab. We punched 2.5 mm diameter matching holes on the PDMS slab and a piece of double-sided tape (Adhesives Research, ARCare 90880) using a 2.5 mm diameter circular punch (Acuderm Inc., P2550). The PDMS slab and the tape were then cut into smaller pieces (∼5 mm on a side) with a razor blade (Sparco, 01485). BIO-133 membranes, PDMS chunks and double-sided tape were further disinfected in 70% ethanol for 2 hours. After disinfection, the BIO-133 membrane was adhered to PDMS using the adhesive tape, so that the matching holes became wells for cell culture. After seeding and growing cells in wells, the assembly was flipped over for diSPIM imaging.

#### Quantification of cell growth

Cured and leached 50 μm thick BIO-133 films were deposited on glass bottom dishes (MatTek, P35G-1.5-14-C). Similar aliquots of U2OS (or HCT 116 TOP1-GFP) cells were seeded onto BIO-133 films or on another glass bottom dish without BIO-133 (MatTek, P35G-1.5-14-C). Dishes seeded with cells were maintained between imaging experiments in an incubator at 37°C, 5% CO_2._ On each dish, a small area was selected and imaged using a widefield microscope equipped with a 10x/0.25 NA objective lens each day, for three days. Cells numbers were estimated from images with the Cell Counter ImageJ plugin (https://imagej.nih.gov/ij/plugins/cell-counter.html). Each experiment was repeated three times. Raw images were divided by Gaussian-blurred versions of themselves (sigma = 5 pixels) to flat-field images prior to display (**Fig. 2a, Supplementary Fig. 2**).

#### Live cell imaging through BIO-133 with diSPIM

U2OS cells were cultured and transfected with 100-200 ng of mEmerald-Tomm20 plasmid (Addgene, 54281) directly on the BIO-133 bottomed well plate. The well plate was inverted and immersed in live cell imaging solution (Invitrogen, A14291DJ). Cells were imaged with a symmetric 0.8/0.8 NA diSPIM, through the BIO-133 layer. 50 volumes were acquired with 3 s intervals between dual-view volumes. Dual-view data were jointly deconvolved with ImageJ plugin DiSPIM Fusion^54^, and were drift-(with ImageJ plugin Correct 3D Drift (https://imagej.net/Correct_3D_Drift) and bleach-corrected (with ImageJ function Bleach Correction (https://imagej.net/Bleach_Correction, exponential fitting method) prior to display.

#### Super-resolution imaging through BIO-133 with iSIM

U2OS or WT HCT-116 cells were cultured and transfected with mEmerald-Tomm20 or LAMP1-EGFP (Taraska Lab, NHLBI) on a 50 μm thick BIO-133 film. The BIO-133 film was cured on a glass bottom dish (MatTek, P35G-1.5-14-C). A 60X, NA = 1.2 water objective (Olympus, PSF grade), correction collar adjusted to 0.17, was used to image the cells through the glass and BIO-133 film using our home built iSIM system^48^ and 488 nm excitation. Volumes were acquired every 3 s for U2OS cells expressing mEmerald-Tomm20 and every 7 s for WT HCT-116 cells expressing LAMP1-EGFP. We used an exposure time of 80 ms, and a z-step of 0.25 μm for U2OS cells and 0.5 μm for WT HCT-116 cells. Live HCT-116 TOP1-GFP cultured on a 50 μm thick BIO-133 film or on a glass bottom dish were imaged to acquire volumes with a step size of 0.5 μm. Raw images were deconvolved with the Richardson-Lucy algorithm for 20 iterations, destriped in Fourier space to remove striping artifacts^66^, and bleach corrected (https://imagej.net/Bleach_Correction). A median filter with kernel size 0.5 pixel was applied to denoise mEmerald-Tomm20 and GFP-LAMP1 images prior to display.

#### Immunolabeling and imaging of multilayered WT HCT-116 cells on BIO-133

WT HCT-116 cells were cultured on a 50 μm thick BIO-133 film on a glass bottom dish until a thick layer was visible by eye. Cells were fixed with 4% paraformaldehyde (Electron Microscopy Sciences) in 1X PBS for 30 minutes at room temperature (RT). Cells were rinsed 3 times in 1X PBS and permeabilized with 0.1% Triton X-100/PBS (Sigma, 93443) for 15 min at RT. Permeabilized cells were rinsed 3 times with 1X PBS and incubated in 1X PBS with primary antibody Rabbit-α-Tomm20 (Abcam, ab186735) and Mouse-α-LaminA/C (Abcam, ab244577) at a concentration of 1:100 for 1 hour at RT. After primary antibody staining, cells were washed in 1X PBS for 5 min, three times. Cells were stained in 1X PBS with secondary antibody Donkey-α-Rabbit Alexa Fluor 488 (Jackson Immuno Research, 711-547-003), Donkey-α-Mouse JF549 (Novusbio, NBP1-75119JF549) and Alexa Fluor 647 Phalloidin (Thermofisher, A22287) at a concentration of 1:100 for 1 hour at RT. Cells were washed in 0.1% Triton X-100/PBS for 5 min, three times. In each spectral channel, 46 slices were acquired on iSIM with an exposure time of 100 ms and a z-step of 0.5 μm. Raw images were deconvolved with the Richardson-Lucy algorithm for 20 iterations and destriped in Fourier space to remove striping artifacts^66^. The 633 nm channel (Alexa Fluor 647 Phalloidin) was bleach corrected (https://imagej.net/Bleach_Correction) across the z stack to compensate for decreased signal further into the stack.

#### Flow cytometry preparation

Sample handling channels, 1 mm wide and 70 μm high, were formed by pouring 20 mL PDMS (Dow Corning, Sylgard 184) on a positive mold made of packaging tape (Duck Brand) cut to the desired dimensions with a craft cutter (Silhouette Cameo) and stuck in a 10 cm Petri dish. A thin PDMS membrane (∼0.5 mm) was air plasma (Harrick Plasma, PDC-32G (115V)) bonded to the channel surface. Holes at the endpoints of the channel were created by punching the PDMS membrane with a 1 mm diameter circular punch (Acuderm Inc., P150) after plasma treatment. A 400 μm wide (142 μm height) imaging channel was cut directly from double-sided silicon-based adhesive tape (Adhesives Research, ARCare 90880) with a craft cutter and stuck to the PDMS device, thus creating a connection between the two sample handling channels in the lower layer. A thin BIO-133 membrane (50 μm) was placed on top of the tape to seal the channel. Two thick PDMS pieces with holes (6 mm and 2 mm diameter) were cut and air plasma bonded to the device to provide fluidic access. When imaging, cells were added to the 6 mm diameter reservoir, and output tubing (Dow Corning, 508-004) was connected to the 2 mm hole. Flow speed was adjusted by changing the height of the output tubing. See also **Supplementary Fig. 5**.

#### Flow cytometry imaging of fixed, DAPI-stained U2OS cells

U2OS cells were fixed in 4% Paraformaldehyde/PBS and subsequently stained with DAPI in 0.1% Triton X-100/PBS (Sigma, 93443). The flow device was mounted on a 10 cm petri dish for imaging with diSPIM. Fixed cells were added to the input port, producing steady flow through the channel after several minutes. 1000 frames of the same image plane were acquired with diSPIM at 50 frames per second under ‘fixed sheet mode’.

### Live animal/tissue imaging through BIO-133 with diSPIM

#### Live imaging of BIO-133 embedded Drosophila wings

A 13 μm thick BIO-133 film was created and cut into two rectangular pieces (4.5 mm x 10 mm) and a square piece (10 mm x 10 mm). The rectangular pieces were deposited on a 10 cm petri dish to form a 1 mm wide open-top channel. Early pupal fly wings were deposited into the channel (convex side up) with 20-40 μL culture media, the square BIO-133 piece placed on top to close the channel and additional culture media was carefully added to the dish (**Fig. 3a, Supplementary Fig. 8)**. Single-view diSPIM imaging was then performed. 360 volumes were acquired with 5 s inter-volume spacing, over 30 minutes. Volumes were deconvolved with MATLAB and bleach corrected with ImageJ (https://imagej.net/Bleach_Correction, exponential fitting method).

#### Fabrication of microfluidics for C. elegans immobilization

Standard soft lithography techniques^67^ were used to fabricate an SU-8 (Kayaku Advanced Materials, formerly Microchem Corp.) master mold for sets of four microfluidic funnels for worm confinement as described^53^. To fabricate devices in BIO-133 (MY Polymers Ltd.), we placed two spacers (100 μm, Precision Brand 44910) beside the pattern, poured polymer onto the mold, covered the mold with a glass slide and cured the polymer under a UV lamp (365nm, Spectroline ENB-280C) for 2 minutes. After curing, we peeled the BIO-133 off the mold, punched inlet and outlet holes with a 1 mm diameter circular punch (Acuderm Inc., P150), and sealed the device to a #1.5 cover glass (24 mm x 50 mm, VWR 48393-241) with double-sided silicone-based adhesive tape (Adhesives Research, ARCare 90880). We cut out an aperture from a 10 cm petri dish and used UV-curable optical cement (Norland Products Inc., Norland Optical Adhesive NOA 68) to secure the coverslip carrying the microfluidic device over the aperture in the petri dish. Inlet and outlet tubing (Dow Corning, 508-004) was clamped to the assembly using a pair of hollow magnets (K&J Magnetics, R211-N52) placed above and below the coverslip, as described^68^. Optical cement was again used to secure tubing to the magnets. See also **Supplementary Fig. 10**.

#### Live imaging of C. elegans through BIO-133 chambers

To load worms into the immobilization device, we added a drop of M9 buffer containing worms to the inlet and created vacuum at the outlet using a syringe. Within several minutes (for a 4-channel chip), worms were observed to align in the channels. The petri dish was then filled with water and worms were imaged with symmetric 0.8/0.8 NA diSPIM. For structural imaging, we added 0.25 mM levamisole to the buffer to stop residual worm motion. For calcium imaging, the outlet was connected to a peristaltic pump (Dolomite Microfluidics, 3200243) which provided negative pressure to immobilize worms without using anesthetics. We simultaneously imaged nuclei structure (TagRFP) and the nuclear-localized calcium response (GCaMP) with 488 nm and 561 nm excitation (Coherent) and image splitting devices on the detection side (Hamamatsu W-VIEW GEMINI), using a previously described fiber-coupled diSPIM system^46^. Dual-view stacks were acquired every 0.8 s over 500 time points. Dual-color, dual-view images were deconvolved and registered with ImageJ plugin DiSPIM Fusion^54^.

#### Droplet-based immobilization of nematodes prior to imaging, and recovery after imaging

*C. elegans* were directly transferred from agar plates into a drop (∼10 μL) of BIO-133 or 10% PEG-DA (ESIBIO GS705) on a #1.5 cover glass (24 mm x 50 mm, VWR, 48393-241). The droplet was positioned between two 100 μm spacers (Precision Brand, 44910), and was compressed by a glass slide followed by 2-minute polymerization under a UV lamp (365nm, Spectroline ENB-280C). After polymerization, worms were immobilized in the resulting gel disk. The gel disk was then placed in a 10 cm petri dish or a standard chamber for diSPIM imaging. Single-view stacks were acquired every 0.25 s for 250 time points. After imaging, worms could be released and checked for viability by gently breaking the droplet with forceps. In some experiments, we immersed the disks in M9 buffer for up to 12 hours, finding that live worms could also be recovered after this period.

#### Tracking nuclei, calcium imaging analysis

TagRFP volumes were imported into Imaris and neurons tracked with Imaris for Tracking (https://imaris.oxinst.com/products/imaris-for-tracking) to obtain the center of each neuron at every timepoint. A custom MATLAB script was used to extract the calcium signal. For every neuron, the average intensity of TagRFP channel I_561_ and the intensity of GCaMP channel I_488_ were computed by averaging pixels within a 2 μm (adult) or 1.5 μm (larval) diameter sphere placed around each center position. I_561_ and I_488_ were calculated from dual-view deconvolved images (Fig. 3 f, g) or single-view raw data (Fig. 3 i, j). The ratio R=I_488_/I_561_ was used to minimize non-GCaMP fluctuations. Neuronal activity for the datasets in **Fig. 3** was reported as dR/R = (R – R_0_)/R_0_, where R_0_ is the baseline for an individual neuron defined as its lower 20th percentile intensity value.

### Chemical and optical perturbations in BIO-133 based imaging devices

#### Fabrication of chemical delivery devices for cells

A modified PDMS well plate design (**Supplementary Fig. 11**) was applied to deliver chemical perturbations to cells (**Fig. 4a-c**). A 400 μm thick BIO-133 film was created using the method described above. 10 mL PDMS was cured in a 10 cm dish, PDMS tubing (Dow Corning, 508-004) was placed on the cured PDMS layer at 8 mm intervals, and another 15 mL PDMS was added to obtain a ∼4 mm thick PDMS slab with channels contained inside. Holes crossing the channels were punched at 8 mm intervals using a 5 mm diameter circular punch (Acuderm Inc., P550). A piece of double-sided silicone-based adhesive tape (Adhesives Research, ARCare 90880) was also punched at 8 mm intervals using a 2.5 mm diameter circular punch (Acuderm Inc., P2550). The PDMS slab and the tape were cut into two-well pieces. BIO-133 film, PDMS chunks and double-sided tape were disinfected in 75% ethanol. After disinfection, the BIO-133 membrane was adhered to PDMS via the double-sided adhesive tape, so that the matching holes became wells for cell culture. After growing cells, tubing (Scientific Commodities Inc., BB31695-PE/2) was inserted into both sides of the channel for introducing chemical flow and another piece of double-sided tape without holes was used to seal the wells. The assembly was flipped over for diSPIM imaging through the BIO-133 membrane.

#### Mitochondrial imaging in the presence of CCCP

U2OS cells were cultured in two 5 mm diameter BIO-133 bottomed wells with ports and transfected with 300-400 ng of mEmerald-Tomm20. Before imaging, the wells were filled with live cell imaging solution (Invitrogen, A14291DJ), flipped over and attached to a 10 cm petri dish with double-sided silicone-based adhesive tape (Adhesives Research, ARCare 90880). For the well containing control cells, tubing was left disconnected from a source. For the well containing cells that experienced chemical perturbation, input tubing was connected to a syringe containing 0.05 mM carbonyl cyanide m-chlorophenyl hydrazine (CCCP, Sigma, C2759). The syringe is higher than the output tube so that drug flow was induced by gravity. We used a valve (McMaster-Carr, 7033T21) placed between the input tube and syringe to control the flow. The valve was closed prior to imaging. For each well, two cells were chosen for imaging. A multi-position acquisition was set in the Micro-manager^69^ diSPIM plugin^70^ to sequentially image the four cells. Volume acquisition time was 3 s, and 90 volumes were acquired for each cell with 60 s intervals between volumes. 10 minutes after the imaging started, the valve was opened and drug flow was induced in ∼60 s. Dual-view images were deconvolved with ImageJ plugin DiSPIM Fusion^54^, drift corrected (ImageJ plugin Correct 3D Drift, https://imagej.net/Correct_3D_Drift) and bleach corrected (ImageJ function Bleach Correction, https://imagej.net/Bleach_Correction, exponential fitting method).

#### Encapsulation of C. elegans into PEG hydrogels

*C. elegans* were encapsulated into PEG hydrogel disks as described in our prior work^7^. PEG hydrogel precursor solutions were prepared by combining 20% w/v poly(ethylene glycol) diacrylate (PEG-DA, 3350 MW, 94.45% acrylation, ESI BIO) with 0.10% w/v Irgacure 2959 photoinitiator (2-hydroxy-4′-(2-hydroxyethoxy)-2-methylpropiophenone, I2959, BASF) in 1x S-basal buffer (100 mM NaCl, 50 mM KPO_4_ buffer, pH 6.0). Clean 24 mm x 50 mm glass coverslips (VWR) were rendered permanently hydrophobic by exposure to vapors of (tridecafluoro-1,1,2,2-tetrahydrooctyl) trichlorosilane (Gelest). For covalent attachment of the PEG hydrogel to glass, coverslips (Thermo Scientific) were silanized by coating with 3-(trimethoxysilyl)propyl methacrylate (TMSPMA, Sigma-Aldrich). Both methods of surface modification were applied to 1” x 3” glass slides (VWR). A small volume (1.75 µL) of PEG hydrogel solution with photoinitiator was pipetted onto a hydrophobic glass slide flanked by two PDMS spacers whose thickness matched the desired hydrogel thickness of 150 μm. Animals were transferred into the hydrogel solution by worm pick. A coverslip, TMSPMA treated for making mounted PEG-DA gels, or untreated for making freestanding gels, was placed over the hydrogel droplet and supported by the PDMS spacers. The glass slide/coverslip sandwich was then placed over a UV light source (312 nm, International Biotechnologies, Inc, model UVH, 12W) and illuminated for two minutes until gelation. Hydrogel disks were immediately transferred to wet agar dishes to keep embedded animals hydrated.

#### Fabrication of microfluidic devices for chemical stimulation of C. elegans

Microfluidic chambers were prepared using poly(dimethyl siloxane) (PDMS; Sylgard 184, Dow Corning) in a ratio of 1:10 and poured to a depth of 5 mm on a silicon master positive mold of the microchannels used previously^71^. Once cut free of the master, devices were punched so that two balanced-length inlets and the outlet had 1.5 mm holes going through the thickness of the material, and 1 mm holes punched from the side to allow flexible tubing to be inserted from the sides. The smooth PDMS device surface opposite the microchannels was irreversibly bonded to a glass slide using oxygen plasma (Harrick PDC-32G, 18W, 45 seconds). A thin PDMS membrane (150 μm) was cut with a 3.5 mm diameter dermal punch and then oxygen plasma bonded to the microfluidic channel surface with the hole in the membrane exposing the micropost array. The hole in the thin PDMS membrane formed a “well” that hydrogel disks could be gently placed in with forceps. A thin BIO-133 membrane (75-80 μm) was prepared by gelation of BIO-133 liquid polymer between two glass slides rendered permanently hydrophobic as described above. Two layers of clear cellophane tape (height of ∼80 μm) formed the standoffs that determined final membrane height. After degassing the microfluidic device in a vacuum chamber for approximately 45 minutes, the device was removed from the desiccator, connected to tubing, and flushed with S. basal buffer before use to remove any air bubbles. Hydrogel disks could be interchanged between the well formed by the PDMS above the micropost array easily using forceps and then the system sealed for microfluidic flow using the thin BIO-133 membrane described above. See also **Supplementary Fig. 12**.

#### Preparation of Chemosensory Stimulus

For both wide-field and diSPIM assays, diacetyl (2,3-butanedione, Sigma) was diluted to 1.1 µM in 1x S. basal (10^−7^ dilution). 1 µL of 1 mg/mL fluorescein solution was added to 40 mL of diacetyl solution to visualize stimulus delivery.

#### Wide-field Imaging with Chemical Stimulation

For wide-field, single plane imaging of multiple *C. elegans* at once^57^, the microfluidic chamber, valves, tubing and reservoirs were prepared as above and placed on a Zeiss AxioObserver epifluorescence microscope with a 5x, 0.25 NA objective, EGFP filter set, and Hamamatsu Orca-Flash 4 sCMOS camera. Micromanager scripts^72^ automatically synchronized capture of ten 30-s trials recording at 10 fps with 10 ms excitation pulses and 10 s chemical stimulation. NeuroTracker software^71^ analyzed the wide-field neural imaging data, from which background-corrected fluorescence changes were calculated in MATLAB as ΔF/F_0_, where F_0_ is baseline neural fluorescence during the four seconds prior to stimulation. Data for multiple individual animals were also presented as a population mean to show the relative decrease in average calcium response after multiple stimulation periods.

#### DiSPIM Imaging with Optical and Chemical Stimulation

Stimulus control for optical illumination or chemical pulses was integrated with diSPIM volumetric imaging using a custom Micromanager script controlling an Arduino Uno and enabling independent digital switching of 6 TTL channels at the beginning of specified image acquisition timepoints. One TTL channel controlled the intensity of a red LED (617 nm, 3W, Mightex) connected to the bottom port of the diSPIM and illuminated the sample through a Nikon 4x, 0.1 NA lower objective. A second TTL channel controlled a 12V fluidic valve system for chemical stimulation (ValveLink 8.2, Automate). Pinch valves allowed flow of either buffer or chemical stimulus lines into the microfluidic channel network, flowing to a common outlet.

For optogenetic stimulation experiments, animals were embedded in BIO-133 disks bonded to a cover glass placed in the diSPIM sample chamber. To embed animals, they were first transiently immobilized by being picked onto seeded (OP50 *E. coli*) plates with 1 mM tetramisole, and allowed to rest for 1.5 hours. Subsequently, worms were picked into a droplet of BIO-133 polymer liquid and gelled in the same manner as the PEG-DA hydrogel disks above, using a TMSPMA silanized coverslip for covalent bonding.

For chemical stimulation experiments, animals were embedded in PEG hydrogel disks. Animals can be maintained in these disks for many hours if they are kept hydrated^7^. Just prior to an experiment, an animal-embedded disk was inserted into the sample cavity of the diSPIM microfluidic chamber. A 75-80 μm thick BIO-133 membrane was sealed to the microfluidic device surface, closing the fluidic channel with the PEG disk and animals contained within. The hydrogel disk was inserted into a droplet of S. basal buffer present in the well to avoid the introduction of bubbles that would disrupt microfluidic flow. To assure continuous microfluidic flow through the chamber without leaking, we balanced inlet and outlet flows by adjusting the reservoir heights. Specifically, inlet reservoir heights were held slightly above the stage (Δ*h*_in_), and the outlet reservoir level was placed further below the stage (Δ*h*_out_> Δ*h*_in_) to ensure a slight negative pressure in the chamber. Microfluidic stimulus switching was achieved using a dual pinch valve (NResearch Inc., 161P091), that alternately allows either a buffer or stimulus line to flow through the microfluidic chamber to the single outflow line.

A typical diSPIM acquisition captured one volume per second (10 ms exposure, minimum slice time setting, 55 slices per volume, 166.4 × 166.4 × 82.5 μm total volume space) for 10 minutes, with 20-s duration stimulation every minute. Z-Projection time series videos were produced in ImageJ from cropped versions of the total number of images, then analyzed for GCaMP neural fluorescence using rectangular boxes for integrated fluorescence density, with a nearby region void of signal used for background subtraction.

