## Supplementary Information for "A polymer gel index-matched to water enables diverse applications in fluorescence microscopy"

**Supplementary Video 1, Mitochondrial dynamics imaged at isotropic resolution with diSPIM, through 50 μm of BIO-133.** U2OS cells expressing mEmerald-Tomm20 were imaged with diSPIM, acquiring 50 volumes with 3 s inter-volume spacing. Lateral and axial maximum intensity projections of dual-view reconstructions are shown. Higher magnification views of red and yellow rectangular regions at left are shown at right. See also **Fig. 2c, d**.

**Supplementary Video 2, Flowing fixed DAPI-labeled U2OS cells, as imaged through 50 μm BIO-133 in diSPIM.** Single-view, raw data are shown. See also **Supplementary Fig. 6.**

**Supplementary Video 3, Mitochondrial dynamics imaged at super-resolution with iSIM, through 50 μm of BIO-133.** U2OS cells expressing mEmerald-Tomm20 were imaged with iSIM, acquiring 25 volumes with 3 s inter-volume spacing. Lateral maximum intensity projections of deconvolved data are shown. A higher magnification view of yellow rectangular regions is also shown. Data have been median filtered for display. See also **Fig. 2f, g.**

**Supplementary Video 4, Lysosomal dynamics imaged at super-resolution with iSIM, through 50 μm of BIO-133.** HCT-116 cells expressing EGFP-LAMP1 were imaged with iSIM, acquiring 60 volumes with 7 s inter-volume spacing. Lateral maximum intensity projections of deconvolved data are shown. Data have been median filtered for display. See also **Supplementary Fig. 7**.

**Supplementary Video 5, Z stack of immunostained Tomm 20, Lamin A/C, and actin obtained with iSIM, through 50 μm of BIO-133.** Multiple layers of HCT-116 cells were grown on a BIO-133 film, fixed, immunostained, and imaged with iSIM. Deconvolved images are shown, with three-color merge shown in lower right images. See also **Fig. 2h, i**.

**Supplementary Video 6, tdTomato-CD4 in *Drosophila* tissue sandwiched between BIO-133 layers**. Single-view diSPIM recordings are shown. 360 volumes were acquired with 5 s inter-volume spacing. The 'red-hot' color map from ImageJ is used for display. See also **Fig. 3b, c.**

**Supplementary Video 7, Pan-neuronal GCaMP6s dynamics imaged at isotropic resolution in immobilized *C. elegans* adults with diSPIM.** Dual-view deconvolved results (GCaMP channel) at 1.25 volumes/s are shown at left, with segmented, tracked dR/R from 126 nuclei shown at right. Lateral and axial views are shown. See also **Fig. 3e-g**.

**Supplementary Video 8, Pan-neuronal GCaMP6s dynamics imaged in immobilized larval *C. elegans*.** Single-view diSPIM recordings at 4 volumes/s are shown at left (GCaMP channel), with segmented, tracked dR/R from 110 nuclei shown at right. Lateral maximum intensity projections are shown. See also **Fig. 3h-j.**

**Supplementary Video 9, Effect of 0.05 mM CCCP on mitochondrial dynamics, as assayed at isotropic resolution with diSPIM, with BIO-133 microfluidics.** U2OS cells expressing mEmerald-Tomm20 were imaged with diSPIM, acquiring 90 volumes with 60 s inter-volume spacing. One BIO-133 well was exposed to 0.05 mM CCCP (left), and another was not (control). Maximum intensity projections of dual-view reconstructions are shown. See also **Fig. 4b, c**.

**Supplementary Video 10, Single-cell GCaMP dynamics under 1.1 μM diacetyl stimulation in immobilized *C. elegans.*** Single-view diSPIM recordings are shown. Lateral maximum intensity projections are shown. See also **Fig. 4f, g.**

**Supplementary Video 11, Single-cell GCaMP dynamics under repetitive optogenetic stimulation in *C. elegans* encapsulated in BIO-133.** Single-view diSPIM recordings are shown. Lateral maximum intensity projections are shown. See also **Fig. 4h, i.**

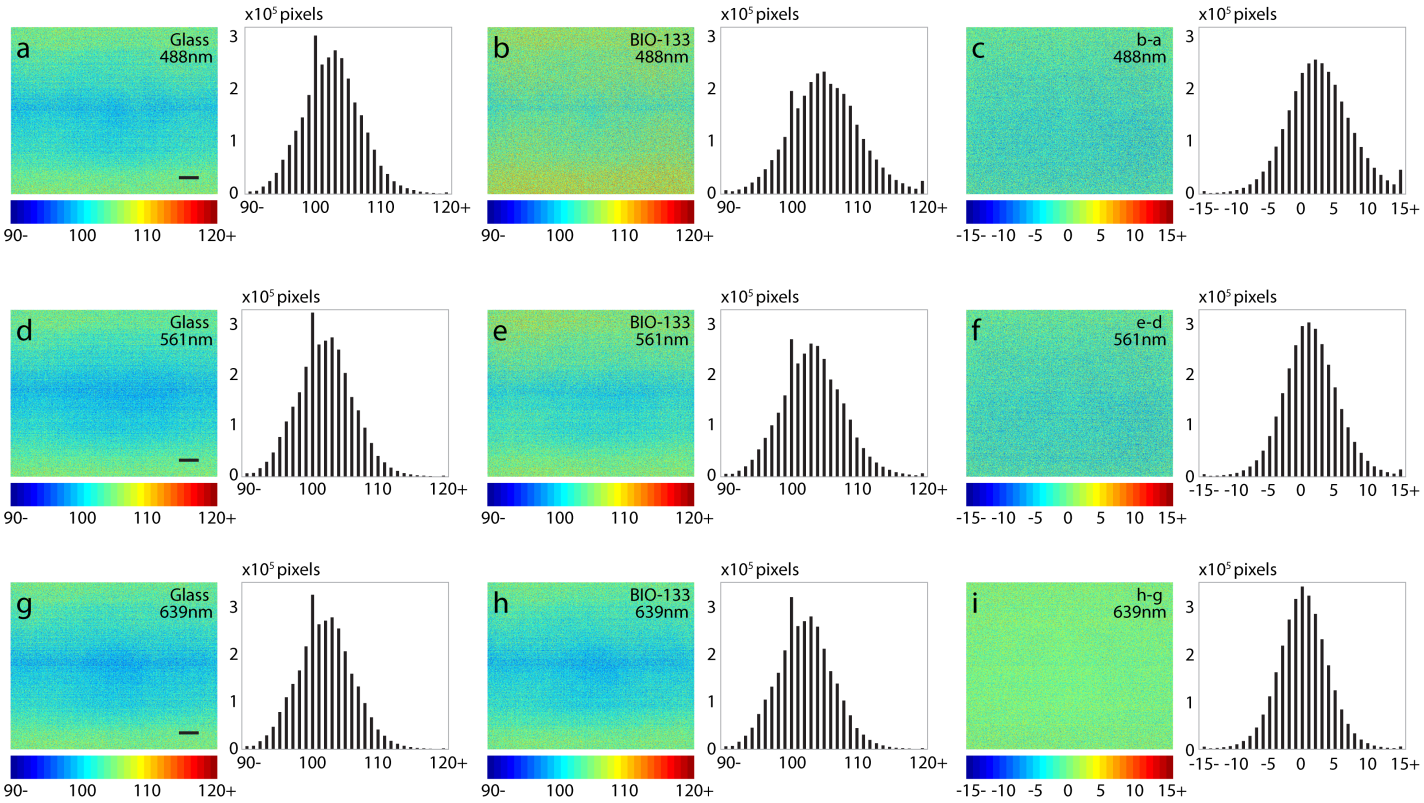


**Supplementary Fig. 1, BIO-133 introduces negligible autofluorescence**. **a)** Image of bare glass coverslip (left) and its associated histogram of counts (right). **b)** As in **a)** but for BIO-133 and its histogram. **c)** Difference image between the two images **b)** - **a)** and associated histogram. Images were acquired with instant SIM using 488 nm excitation and 45 mW excitation. **d-f)** As in **a-c)** but using 70 mW 561 nm excitation. **g-i**) As in **a-c)** but using 90 mW 647 nm excitation. Scale bars: 10 μm.

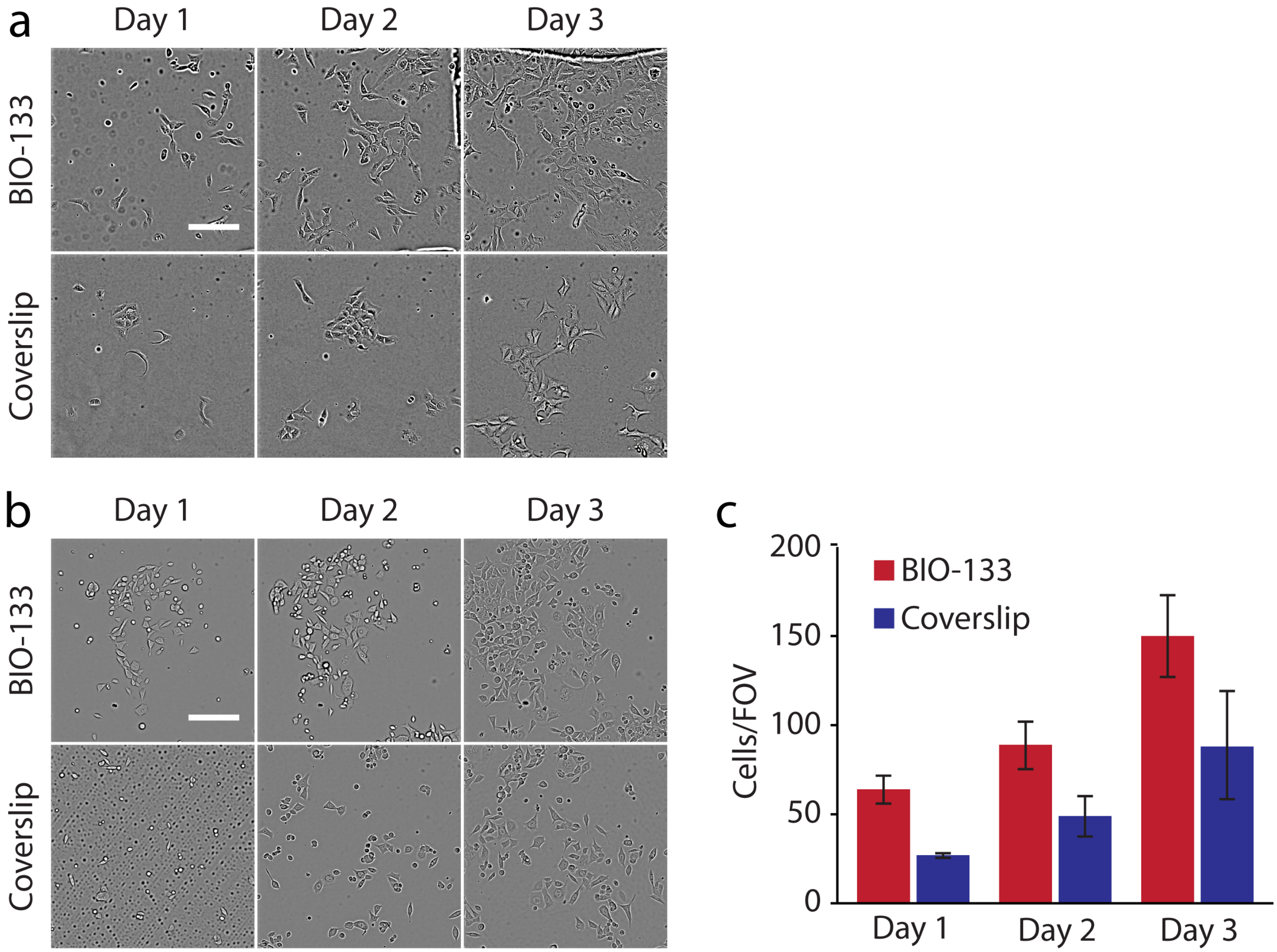


**Supplementary Fig. 2, Cells grown on BIO-133 exhibit similar morphology and growth rate to cells grown on glass coverslips. a)** U2OS cell growth over 3 days, on BIO-133 (top row) and glass (bottom row). **b)** As in **a)**, but for HCT-116 TOP1-GFP cells. **c)** Quantifying HCT-116 TOP1-GFP cell growth on 50 μm thick BIO-133 layer vs. glass coverslip. Means and standard deviations from 3 fields of view are shown. Brightfield images are shown after flat-fielding. See also **Fig. 2a**. Scale bar: 200 μm.

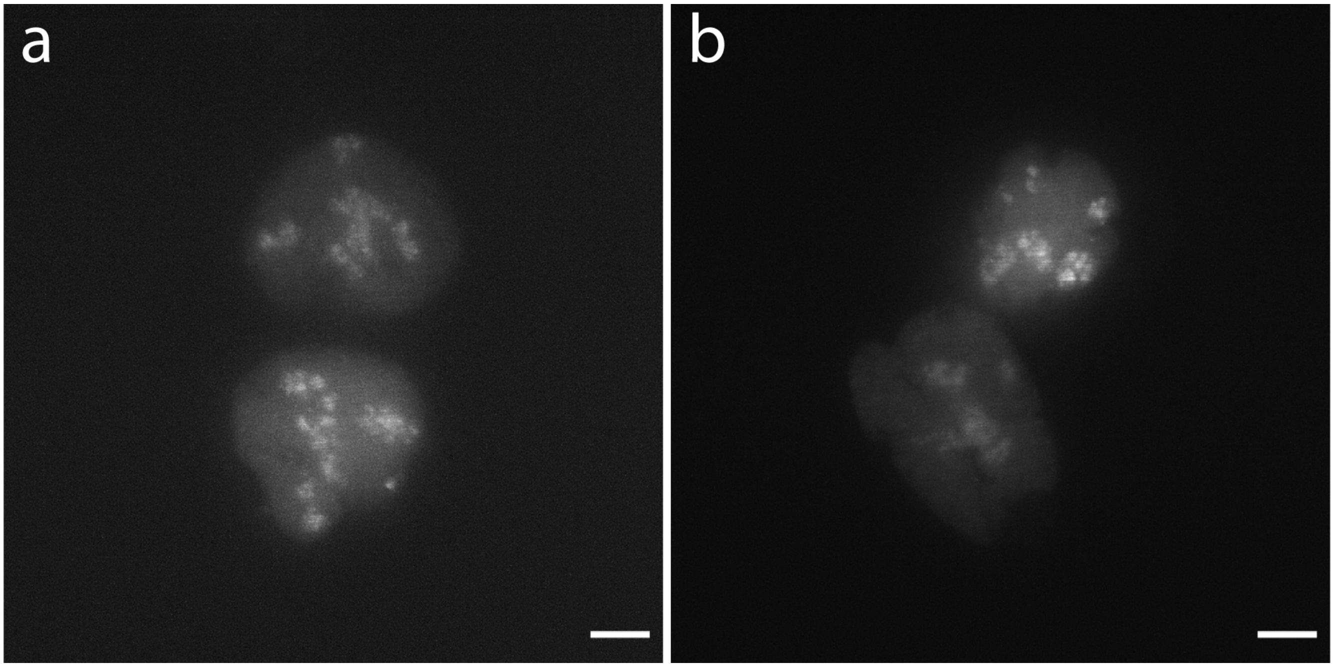


**Supplementary Fig. 3, Endogenous expression and localization of topoisomerase I-GFP in HCT-116 TOP1-GFP cells cultured on BIO-133 is comparable to cells grown on glass coverslips.** Deconvolved maximum intensity projection of iSIM volume showing endogenous expression and localization of topoisomerase I-GFP in HCT-116 TOP1-GFP cells cultured on glass surface **a)** or BIO-133 surface **b)**. Scale bar: 5 μm.

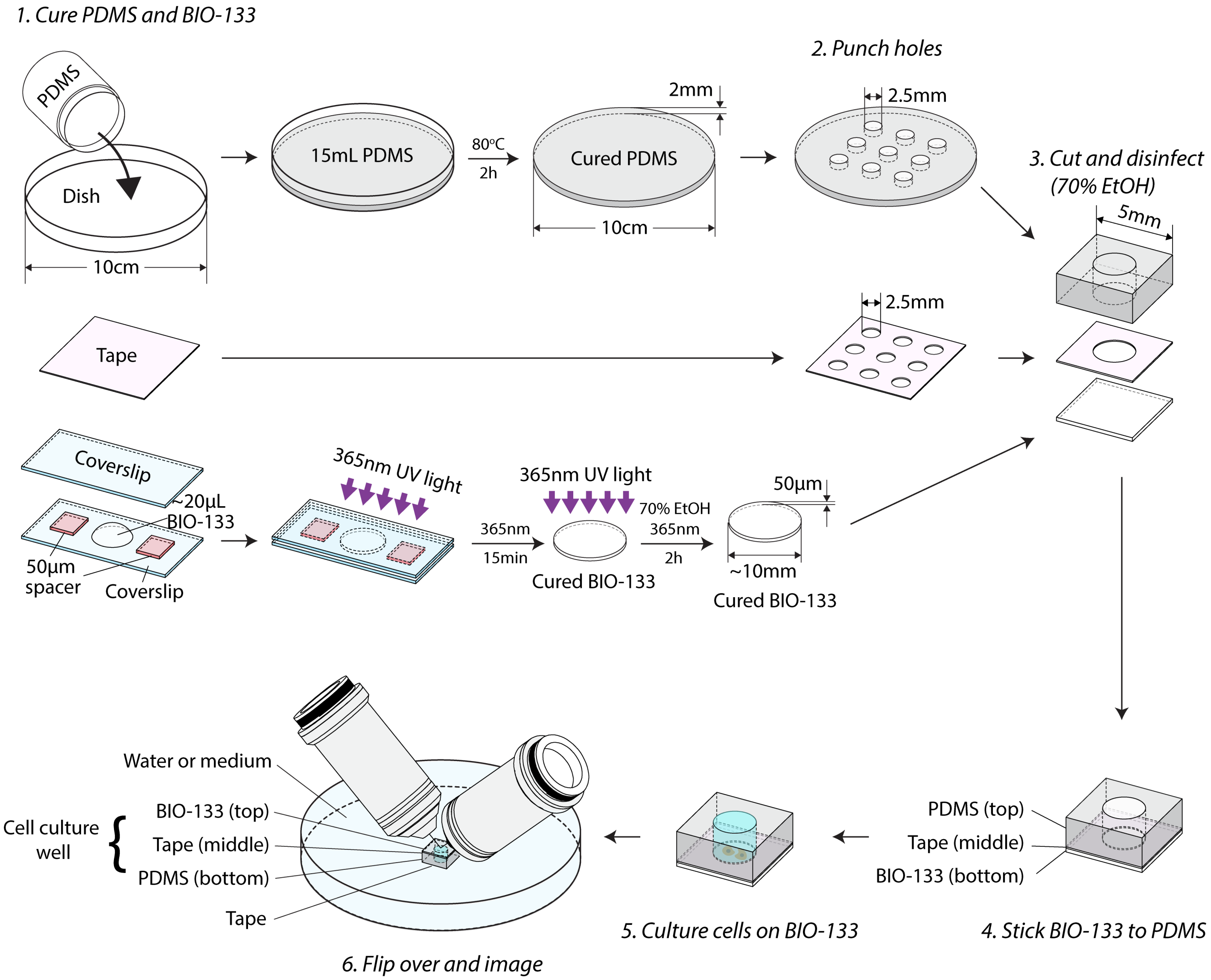


**Supplementary Fig. 4, Fabricating BIO-133-sided substrates for cell culture and diSPIM imaging**. 1) Cure a PDMS slab and a thin BIO-133 film. 2) Punch wells into PDMS and punch holes on a double-sided tape. 3) Cut PDMS, Tape and BIO-133 into desired shape, disinfect. 4) BIO-133 is adhered to PDMS via adhesive tape. 5) Cells are seeded and cultured on BIO-133 film. 6) The assembly is flipped over and imaged in diSPIM.

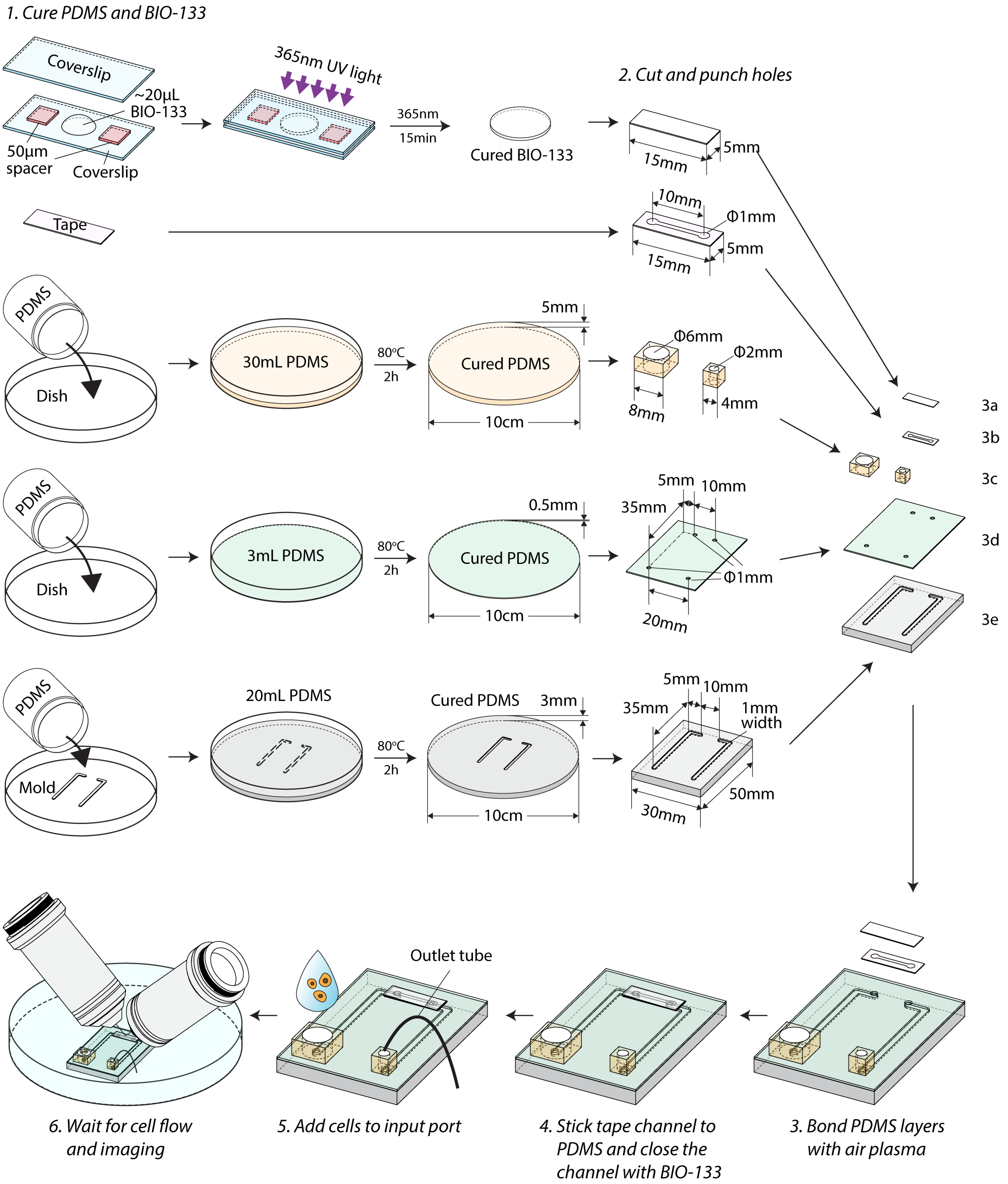


**Supplementary Fig. 5, Flow cytometry with BIO-133**. 1) Cure BIO-133, a PDMS slab, a thin layer of PDMS and a PDMS with channels. 2) Cut to desired shape and punch holes. Note that holes at the ends of the PDMS channel can also be punched after plasma treatment. 3) Bond the three PDMS layers with air plasma (3c, 3d, 3e). 4) Stick tape channel (3b) to PDMS to connect the two PDMS channels. Stick BIO-133 (3a) to tape to close the channel. 5) Add cells to input port. 6) Wait for the cells to flow into the tape channel and image.

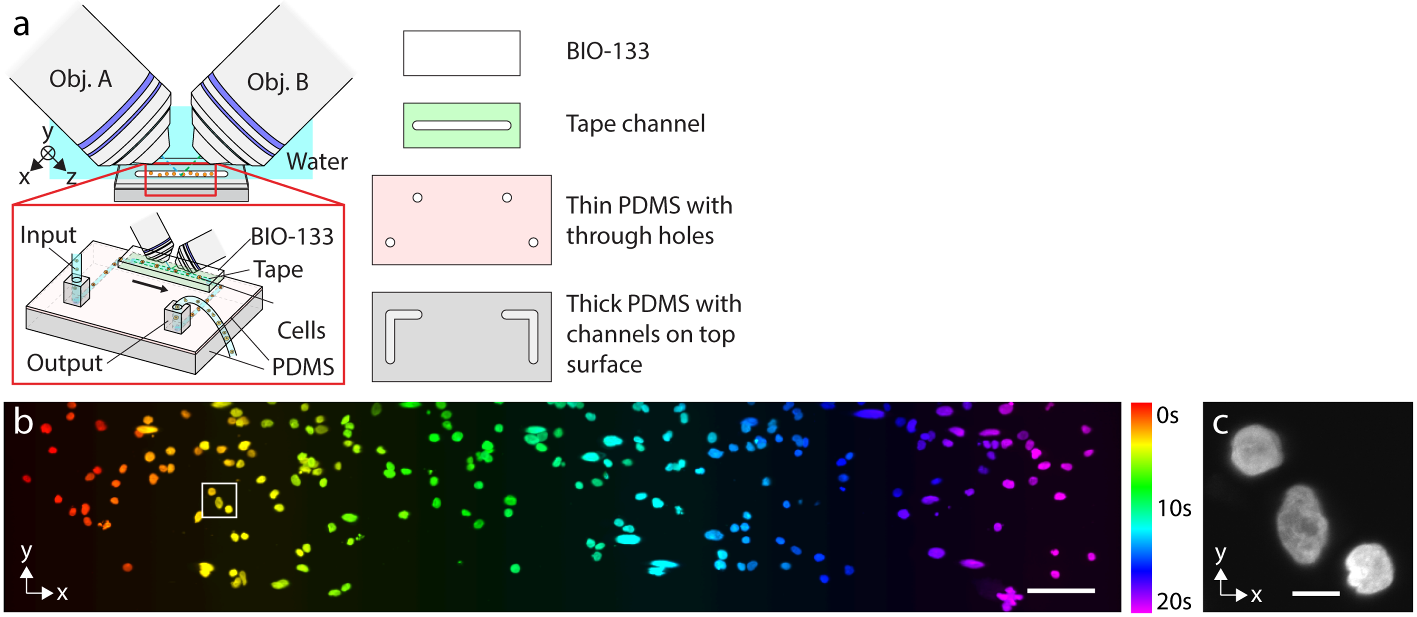


**Supplementary Fig. 6, Simple flow cytometry with BIO-133. a)** Gravity-driven flow channel with BIO-133 top layer, see also **Methods**. **b)** DAPI-stained nuclei in fixed U2OS cells. Multiple fields of view from 20-s recording are stitched together to show flow vs. time. See also **Supplementary Video 5**. Scale bar: 100 µm. **c)** Higher magnification view of nuclei in white rectangular region in **b)**. Scale bar: 10 µm.

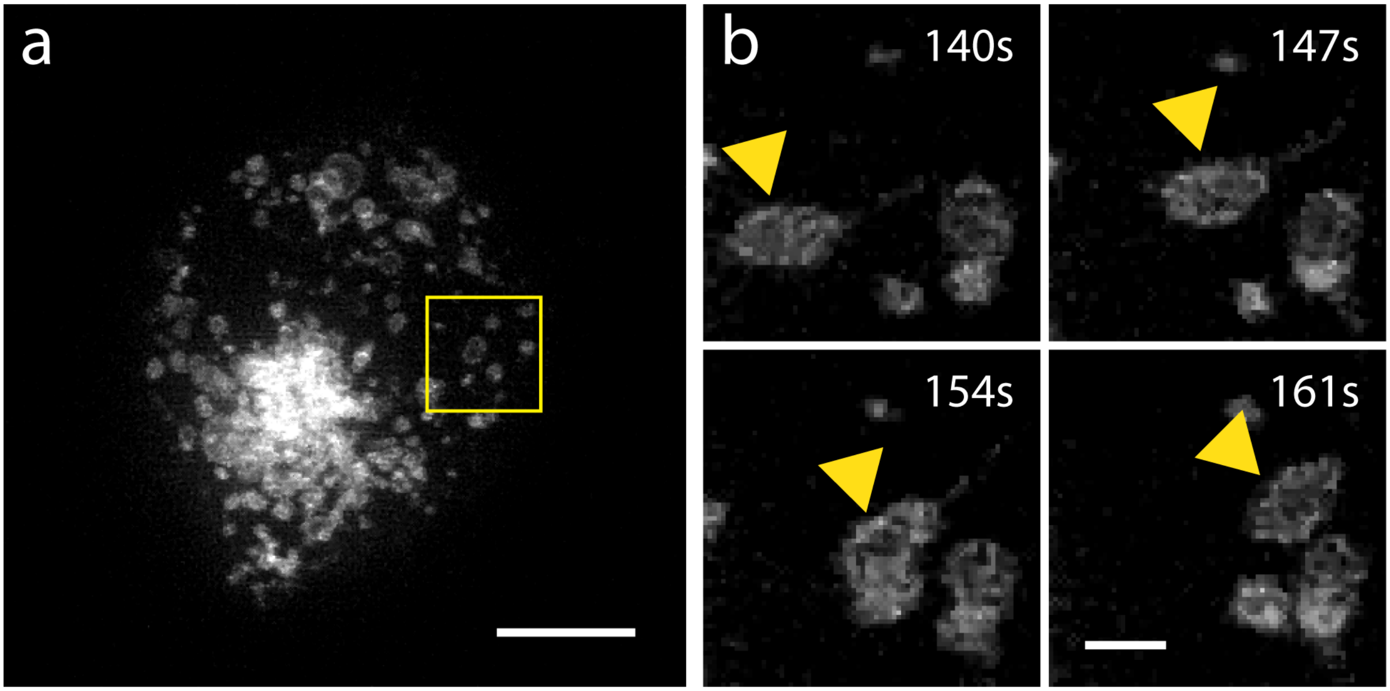


**Supplementary Fig. 7, Lysosome dynamics as imaged via iSIM through 50 μm BIO-133. a)** Deconvolved maximum intensity projection of iSIM volume showing WT HCT-116 cells expressing EGFP-LAMP1. Scale bar: 5 μm. **b)** Higher magnification view of yellow square rectangular region in **a)**, projected over axial region 9-12 μm from the coverslip. Yellow arrowhead marks the same lysosome. See also **Supplementary Video 4**. Median-filtered data (kernel 0.5 pixel) are shown. Scale bar: 1 μm.

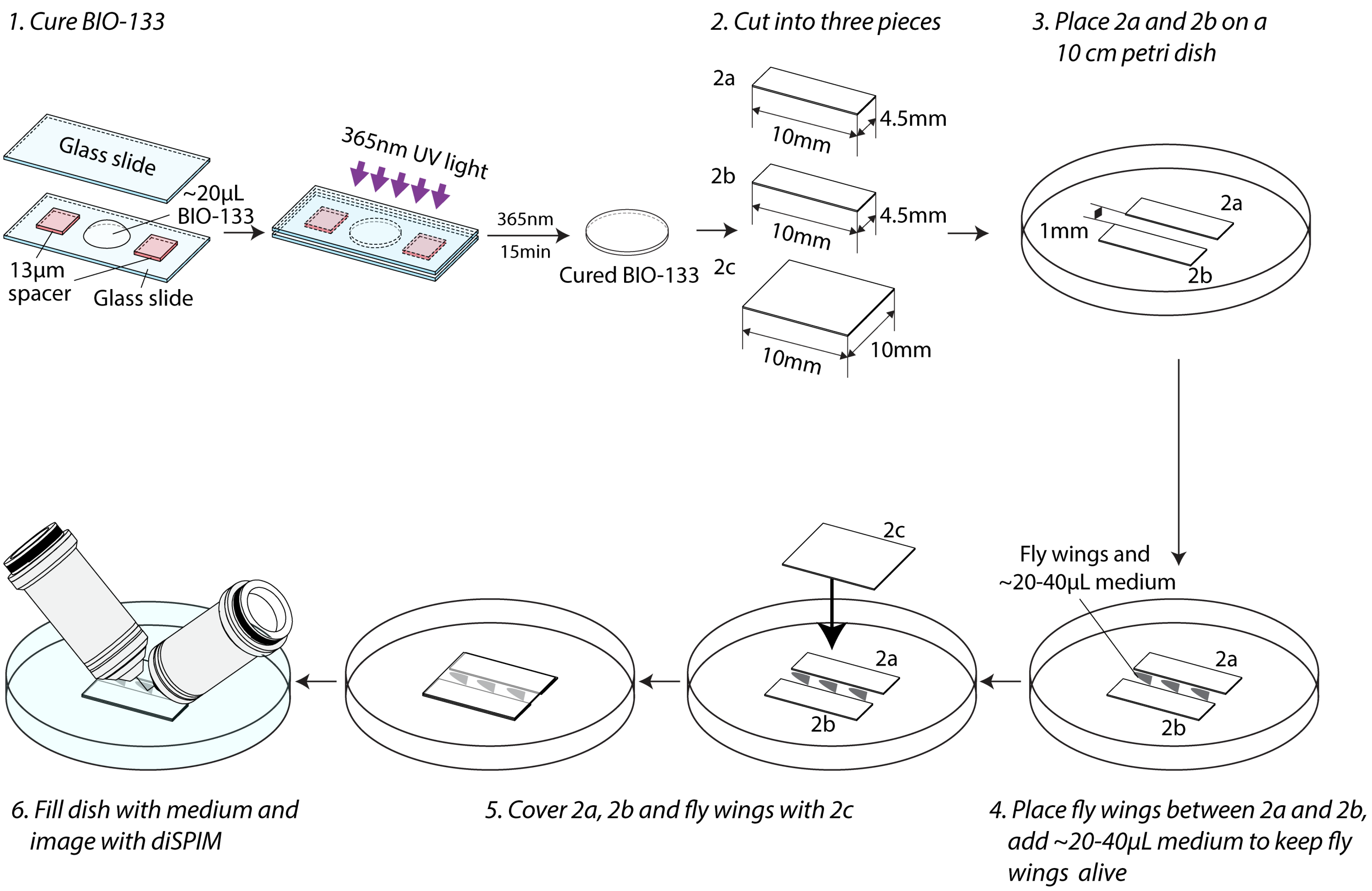


**Supplementary Fig. 8, Imaging fly wings in BIO-133 channel with diSPIM.** 1) Cure 13 µm thick BIO-133 film. 2) Cut into three pieces (2a, 2b and 2c). 3) Place 2a and 2b directly on a 10 cm petri dish to form a 1 mm wide open-top channel. 4) Place the fly wings between 2a and 2b, add ~20 - 40 µL medium. 5) Cover 2a, 2b and fly wings with 2c. 6) Carefully fill the dish with medium and image with diSPIM.

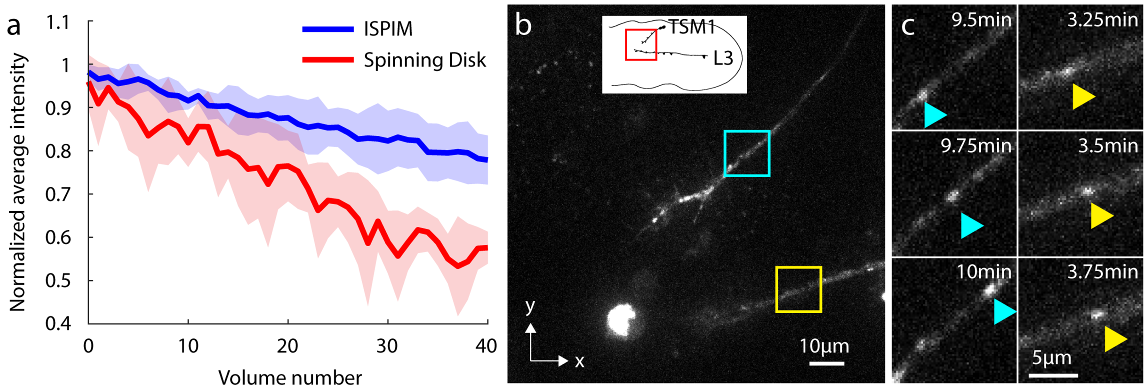


**Supplementary Fig. 9, Imaging fly wings with spinning disk confocal microscopy.** a) Bleaching comparison between iSPIM (**Fig. 3b**) and spinning disk confocal microscopy. iSPIM volumes were acquired every 5 s with 1 µm axial spacing; spinning disk confocal volumes were acquired every 15 s with 1.3 µm axial spacing. Shaded areas encompass one standard deviation around means (curves); data are pooled from 5 different regions in each dataset. b) Example maximum intensity projection from spinning-disk dataset, indicating TSM1 and L3, labeled with tdTomato-CD4. c) Higher magnification views of blue and yellow rectangles in b), emphasizing trafficking CD4 puncta.

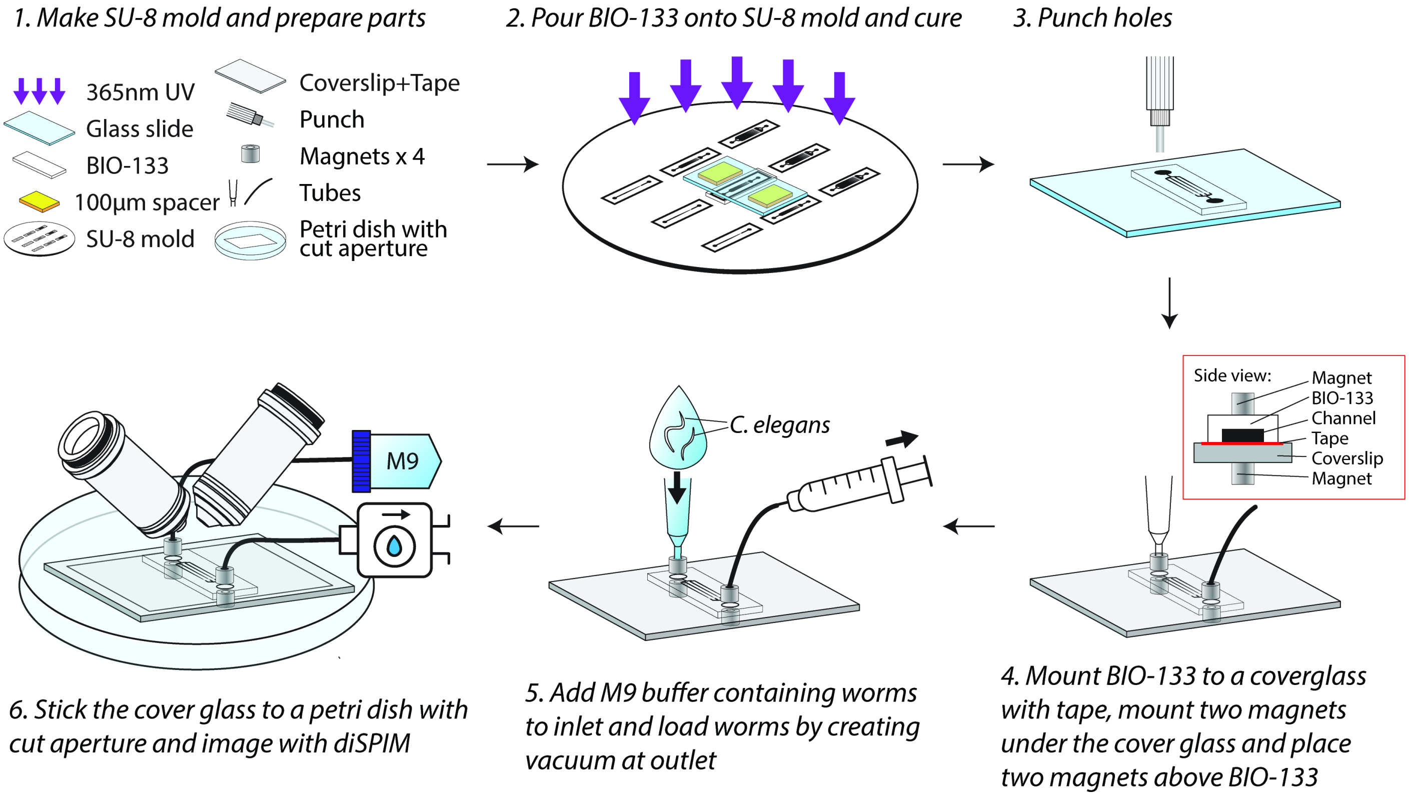


**Supplementary Fig. 10, Immobilization and imaging of *C. elegans* in BIO-133 microfluidics**. 1) The SU-8 mold and other components are assembled. 2) BIO-133 is poured into the SU-8 mold and cured. 3) The BIO-133 mold is peeled off and holes punched at each end. 4) BIO-133 is mounted to a coverglass with adhesive tape, additionally mounting magnets with tubing for fluid delivery/removal. 5) M9 buffer, worms are added to channels. 6) The assembled microfluidic is placed within a 10 cm petri dish with aperture cut out, covered in water, and imaged with diSPIM.

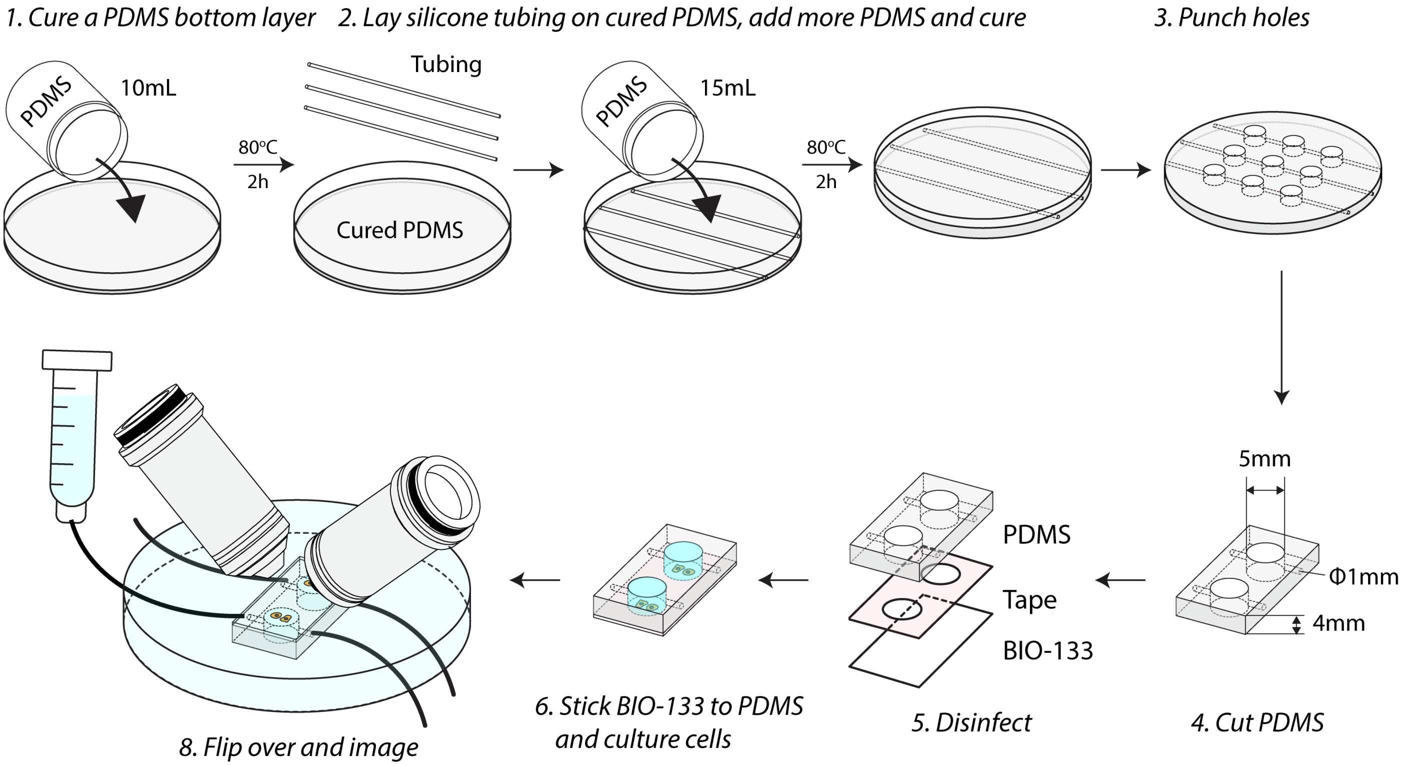


**Supplementary Fig. 11, Fabricating BIO-133-sided substrates for chemically stimulating cells.** 1) Cure a thin layer of PDMS. 2) Lay silicone tubing on the cured PDMS layer, add more PDMS and cure again to create channels. These channels are used to deliver buffer flow or chemical stimulation and are optional. 3) Wells are punched into PDMS. 4) Cut PDMS into desired shape. 5) Cure BIO-133 film, punch holes on double-sided tape, and put all components in 70% EtOH for disinfection. 6) BIO-133 is adhered to PDMS via adhesive tape; cells are seeded and cultured on BIO-133 film. 7) The assembly is flipped over and imaged in diSPIM. For chemical stimulation, tubing is inserted into PDMS and chemical flow is driven by gravity.

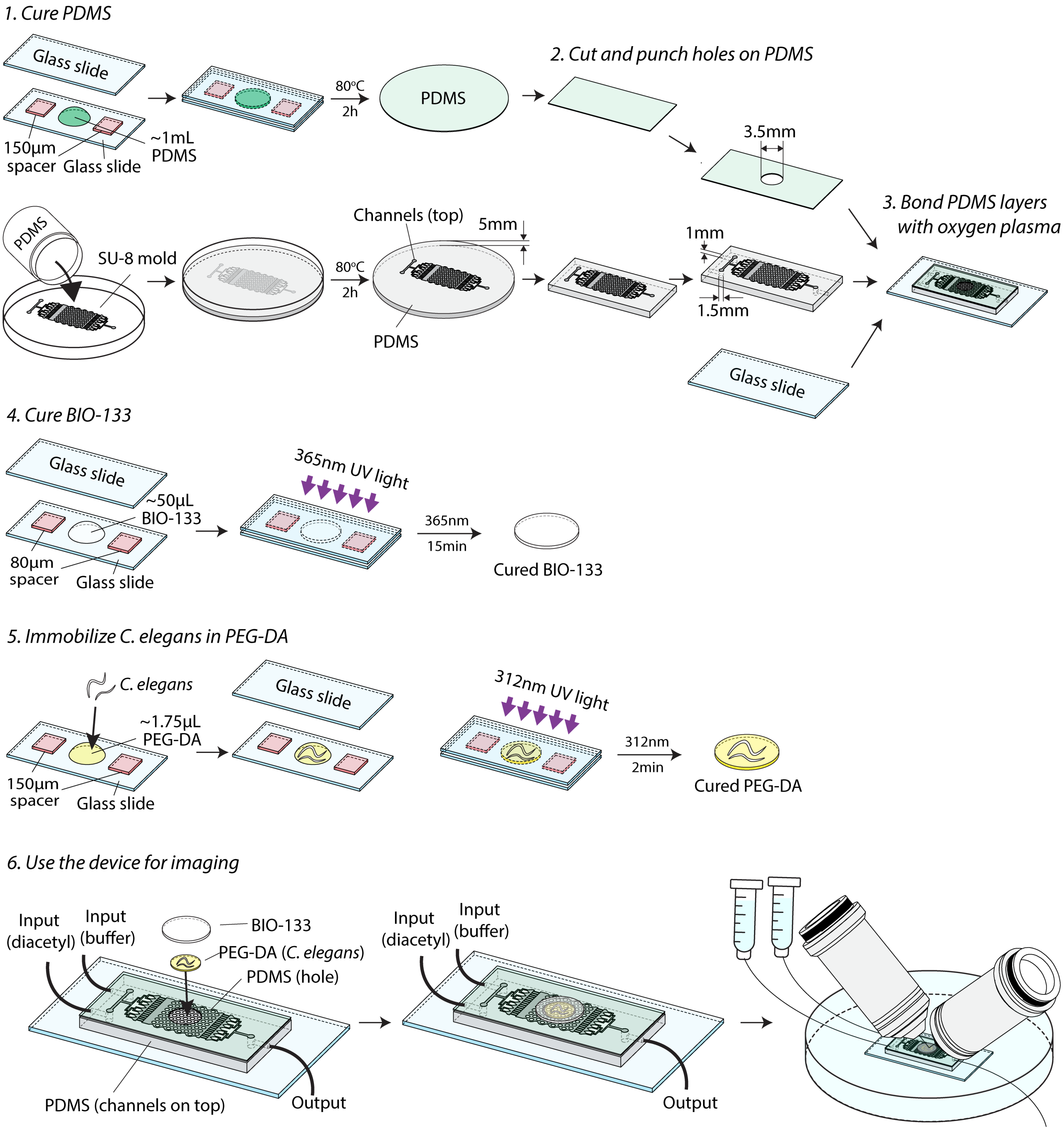


**Supplementary Fig. 12, Fabricating BIO-133-sided substrates for chemically stimulating *C. elegans*.** 1) Make PDMS channels and a 150 μm thick PDMS film. 2) Cut PDMS to desired size. Punch a 3.5 mm diameter hole in the film. Punch input and output ports to the PDMS channel. 3) Bond PDMS layers and glass with oxygen plasma treatment. 4) Cure a 80 μm thick BIO-133 film. 5) Immobilize worms in PEG-DA hydrogel. 6) Put the PEG-DA in the 3.5 mm hole contained within the PDMS film, cover the hole with BIO-133 and image.

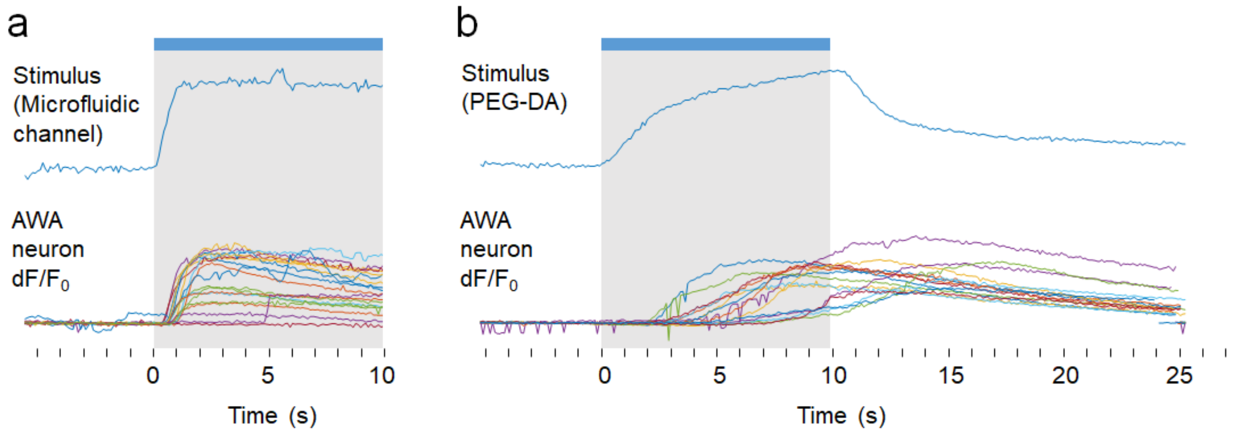


**Supplementary Fig. 13, Diffusion of the chemical stimulus through the BIO-133/PEG-DA/PDMS device delays neural responses by a few seconds.** a) In a microfluidic channel, stimulus (100 ng/mL fluorescein, 1.1 μm diacetyl) switch within 1 s and sensory neurons of animals in the channel respond with a 0.5-1 s delay. Mean fluorescence of fluorescein dye “stimulus” (above) and normalized AWA neuron fluorescence (below). Individual animal responses are shown by colors. b) Animals embedded in a PEG hydrogel disk above a microfluidic channel experience a slower stimulus onset and offset, and a corresponding delay in neural response, due to chemical diffusion into and out of the hydrogel.

**Supplementary Table 1, Apparent size of 100 nm yellow-green beads as imaged through different thicknesses of different polymers**. Full width at half maximum (FWHM) for xyz dimensions as defined as in **Fig. 1c**. Note that defining 'z' this way leads to an underestimation of the FWHM along the direction of greatest elongation for the asymmetric images produced with PEG-DA, PDMS, and FEP. Means and standard deviations are shown; number of measurements is listed in parentheses after each x measurement.

| **FWHM-x(nm)** | | | | |
|---|---|---|---|---|
| Thickness(μm) | BIO-133 | PEG-DA | PDMS | FEP |
| 0 | 395.9±7.7 (70) | 395.9±7.7 (70) | 395.9±7.7 (70) | 395.9±7.7 (70) |
| 25 | 396.5±7.3 (68) | 456.4±15.5 (53) | 816.8±24.9 (48) | 670.5±26.3 (53) |
| 50 | 397.5±7.8 (60) | 512.9±18.0 (42) | 1005.5±64.3 (36) | 734.6±40.4 (76) |
| 75 | 403.8±8.5 (65) | 622.9±22.0 (35) | 1154.0±54.5 (69) | 783.5±46.6 (65) |
| 100 | 406.7±6.4 (70) | 703.1±28.0 (23) | 1235.6±72.4 (55) | |
| 125 | 410.8±9.1 (68) | 736.0±30.7 (42) | 1302.2±56.9 (42) | 1079.2±51.9 (71) |
| 150 | 416.5±8.5 (57) | 816.3±23.1 (38) | 1383.8±53.2 (38) | |
| **FWHM-y(nm)** | | | | |
| Thickness(μm) | BIO-133 | PEG-DA | PDMS | FEP |
| 0 | 400.8±7.6 | 400.8±7.6 | 400.8±7.6 | 400.8±7.6 |
| 25 | 395.2±7.5 | 399.5±6.5 | 433.9±12.6 | 398.8±9.3 |
| 50 | 402.5±10.9 | 410.7±9.0 | 450.8±20.3 | 441.0±12.1 |
| 75 | 399.9±9.5 | 422.7±13.4 | 460.9±22.1 | 478.2±18.1 |
| 100 | 396.1±8.7 | 424.7±14.3 | 483.8±20.5 | |
| 125 | 403.4±12.2 | 417.9±9.6 | 495.5±22.7 | 542.2±25.2 |
| 150 | 408.2±11.4 | 440.7±12.6 | 510.6±29.1 | |
| **FWHM-z(nm)** | | | | |
| Thickness(μm) | BIO-133 | PEG-DA | PDMS | FEP |
| 0 | 1527.9±119.5 | 1527.9±119.5 | 1527.9±119.5 | 1527.9±119.5 |
| 25 | 1527.8±120.5 | 1480.6±88.7 | 1535.4±140.7 | 1603.9±61.6 |
| 50 | 1544.9±133.6 | 1578.5±90.5 | 1550.6±81.1 | 1640.4±73.4 |
| 75 | 1497.8±110.3 | 1397.5±51.7 | 1678.2±102.4 | 1661.2±97.0 |
| 100 | 1542.1±126.5 | 1394.5±37.5 | 1850.1±96.1 | |
| 125 | 1505.9±114.2 | 1445.3±75.8 | 2018.7±127.3 | 1920.6±112.3 |
| 150 | 1436.9±65.6 | 1413.2±48.3 | 2008.3±134.2 | |

**Supplementary Table 2a, Acquisition parameters for all cellular data acquired in this work.** See also **Methods**. Here 'iSPIM' refers to single-view diSPIM.

| **Samples** | | **Live U2OS, mitochondrial label** | | | **Live WT HCT-116, lysosome label** | **Live HCT-116 TOP1-GFP** | **Fixed WT HCT-116, immunolabel** | **Fixed U2OS, DAPI** |
|---|---|---|---|---|---|---|---|---|
| **Figures/Videos** | | Fig. 2 c, d Sup. Video 1 | Fig. 2 f, g Sup. Video 3 | Fig. 4 b, c Sup. Video 9 | Sup. Fig. 7 Sup. Video 4 | Sup. Fig. 3 | Fig. 2 h, i Sup. Video 5 | Sup. Fig. 6 Sup. Video 2 |
| **Fluorescence label** | | mEmerald-Tomm20 | | | LAMP1-GFP | Topoisomerase I-GFP | Lamin A-JF549 Tomm20-AF488 Phalloidin AF647 | DAPI |
| **Microscope** | | diSPIM | iSIM | diSPIM | iSIM | iSIM | iSIM | diSPIM (iSPIM) |
| **View number** | | 2 | 1 | 2 | 1 | 1 | 1 | 1 |
| **Color number** | | 1 | 1 | 1 | 1 | 1 | 3 | 1 |
| **Acquisition** | Excitation, nm | 488 | 488 | 488 | 488 | 488 | 488/561/633 | 405 |
| | Step size x Slices per view per color | 0.5 µm x 150 slices | 0.25 µm x 8 slices | 1 µm x 60 slices | 0.5 µm x 26 slices | 0.5 µm x 24 slices | 0.5 µm x 46 slices | 1 slice |
| | Acquisition time per time point | 2.8 s | 1 s | 3 s | 3 s | - | - | 20 ms |
| | Time interval | 3 s | 3 s | 60 s | 7 s | - | - | 20 ms |
| | Total time points | 50 | 25 | 90 | 60 | 1 | 1 | 1000 |
| | Total acquisition time | 150 s | 75 s | 90 min | 420 s | 3 s | 15 s | 20 s |
| **Data processing** | Registration | √ | x | √ | x | x | x | x |
| | Deconvolution | √ | √ | √ | √ | √ | √ | x |
| | Bleach correction | √ | √ | √ | √ | x | 488x/561x/633 √ | x |
| | Drift correction | √ | x | √ | x | x | x | x |

**Supplementary Table 2b, Acquisition parameters for all tissue/animal data used in this work.** See also **Methods**. Here 'iSPIM' refers to single-view diSPIM and 'PHD' to pleckstrin homology domain, localizing GFP to the cell surface.

| **Samples** | | **Live fly wing** | ***C. elegans* neuron** | ***C. elegans* neuron** | | ***C. elegans* AWA neuron** | | |
|---|---|---|---|---|---|---|---|---|
| **Figures/Videos** | | Fig. 3 b, c Sup. Video 6 | Fig. 3 d | Fig. 3 e Sup. Video 7 | Fig. 3 h Sup. Video 8 | Fig. 4 e | Fig. 4 f Sup. Video 10 | Fig. 4 h Sup. Video 11 |
| **Fluorescence label** | | CD4-tdTomato | PHD-GFP | GCaMP6s-pan-neuronal nuclei, tagRFP-pan-neuronal-nuclei | | GCaMP2.2b | | |
| **Microscope** | | diSPIM (iSPIM) | diSPIM | diSPIM | diSPIM (iSPIM) | Widefield | diSPIM (iSPIM) | diSPIM (iSPIM) |
| **View number** | | 1 | 2 | 2 | 1 | 1 | 1 | 1 |
| **Color number** | | 1 | 1 | 2 | 2 | 1 | 1 | 1 |
| **Acquisition** | Excitation, nm | 561 | 488 | 488/561 | 488/561 | 488 | 488 | 488 |
| | Step size x Slices per view per color | 1 µm x 70 slices | 1 µm x 50 slices | 1 µm x 40 slices | 1 µm x 28 slices | 1 slice | 1.5 µm x 20 slices | 1.5 µm x 30 slices |
| | Acquisition time per time point | 3.8 s | - | 0.7 s | 245 ms | 10 ms | 0.56 s | 0.68 s |
| | Time interval | 5 s | - | 0.8 s | 0.25 s | 100 ms | 1 s | 1s |
| | Total time points | 360 | 1 | 450 | 250 | 300 | 45 | 600 |
| | Total acquisition time | 30 min | 2.3 s | 360 s | 62.5 s | 30 s | 45 s | 10 min |
| **Data processing** | Registration | x | √ | √ | x | x | x | x |
| | Deconvolution | √ | √ | √ | √ | x | x | x |
| | Bleach correction | √ | x | x | x | x | x | x |
| | Drift correction | x | x | x | x | x | x | x |
